# The ribosome synchronizes folding and assembly to promote oligomeric protein biogenesis

**DOI:** 10.1101/2025.05.27.656346

**Authors:** Alžběta Roeselová, Santosh Shivakumaraswamy, Gabija Jurkeviciute, Jessica Zhiyun He, Josef Auburger, Jaro L. Schmitt, Günter Kramer, Bernd Bukau, Radoslav I. Enchev, David Balchin

**Affiliations:** Protein Biogenesis Laboratory, The Francis Crick Institute; London, UK; Visual Biochemistry Laboratory, The Francis Crick Institute; London, UK; Center for Molecular Biology of Heidelberg University (ZMBH), DKFZ-ZMBH Alliance, Heidelberg, Germany; Bayer AG, Research & Development, Pharmaceuticals, D-42113 Wuppertal, Germany; Division of Biosciences, Faculty of Life Science, University College London, United Kingdom; Department of Neuroscience, Physiology and Pharmacology, University College London, United Kingdom; Department of Chemistry, Imperial College London, London, UK

## Abstract

Natural proteins are structurally diverse and often form intricate multidomain, oligomeric architectures. This presents a *prima facie* challenge to cellular homeostasis, as topologically complex proteins seldom refold efficiently in vitro. How cells overcome sequence-intrinsic folding limitations to optimize protein biogenesis is incompletely understood. Here, we show that efficient folding and assembly of the model five-domain homotetramer β-galactosidase is obligatorily coupled to its synthesis on the ribosome, and define the underlying mechanisms. During refolding of full-length protein from denaturant, maturation of the catalytic domain is frustrated. Assembly outpaces monomer folding, and non-native oligomers accumulate. The ribosome directs the order of folding events and specifies the pathway of oligomer assembly. Efficient *de novo* folding is characterised by segmental domain folding, shaped by binding of a nascent amphipathic helix to a cryptic pocket on the ribosome surface. Homomer assembly initiates cotranslationally via recruitment of a full-length subunit to the nascent polypeptide, and the failure to do so results in misassembly. Our results reveal how the ribosome can dictate the timing of folding and assembly to enable efficient biogenesis of a topologically complex protein.

## INTRODUCTION

Small single domain proteins typically refold rapidly and reversibly in vitro, implying that their amino acid sequence suffices to encode productive folding. The “average” protein, however, is large^1^, multidomain^2^ and oligomeric^3^, and refolds slowly or inefficiently in isolation^4^. Based on the relationship between protein length and in vitro refolding kinetics, >600 bacterial proteins are predicted to refold slower than the doubling time of *E. coli*^5^. This discrepancy suggests that the cellular environment contributes additional information to folding reactions of topologically complex proteins.

Folding assistance is provided in part by molecular chaperones, which suppress off-pathway aggregation and catalyse refolding following proteotoxic stress^6,7^. In addition, the translating ribosome may direct efficient *de novo* folding during protein synthesis. The ribosome influences protein maturation at different levels. Proximity to the ribosome surface often alters the stability of nascent domains^8–14^, and the vectorial character of protein synthesis can avoid interdomain misfolding^1,15–17^. For oligomeric proteins, assembly frequently initiates on the ribosome^18–22^. Cotranslational assembly can protect unstable subunits from aggregation^21,23^ and may contribute to the fidelity of protein complex formation^24,25^. The ribosome therefore constitutes a unique folding environment, but how exactly this benefits difficult-to-fold proteins is unclear^26^.

Here, we used *Escherichia coli* β-galactosidase (β-gal), a large homotetramer of five-domain subunits^27^, as a model for understanding how the cellular environment optimises protein folding and assembly (Fig. 1a). We show that maturation of β-gal is highly efficient only when folding is coupled to synthesis on the ribosome, and use structural and proteomic approaches to provide a molecular rationale. We find that the ribosome directs monomer folding and enforces directional assembly of β-gal subunits. The resulting synchronisation between cotranslational folding and assembly avoids kinetic traps that frustrate refolding.

**Fig. 1.**
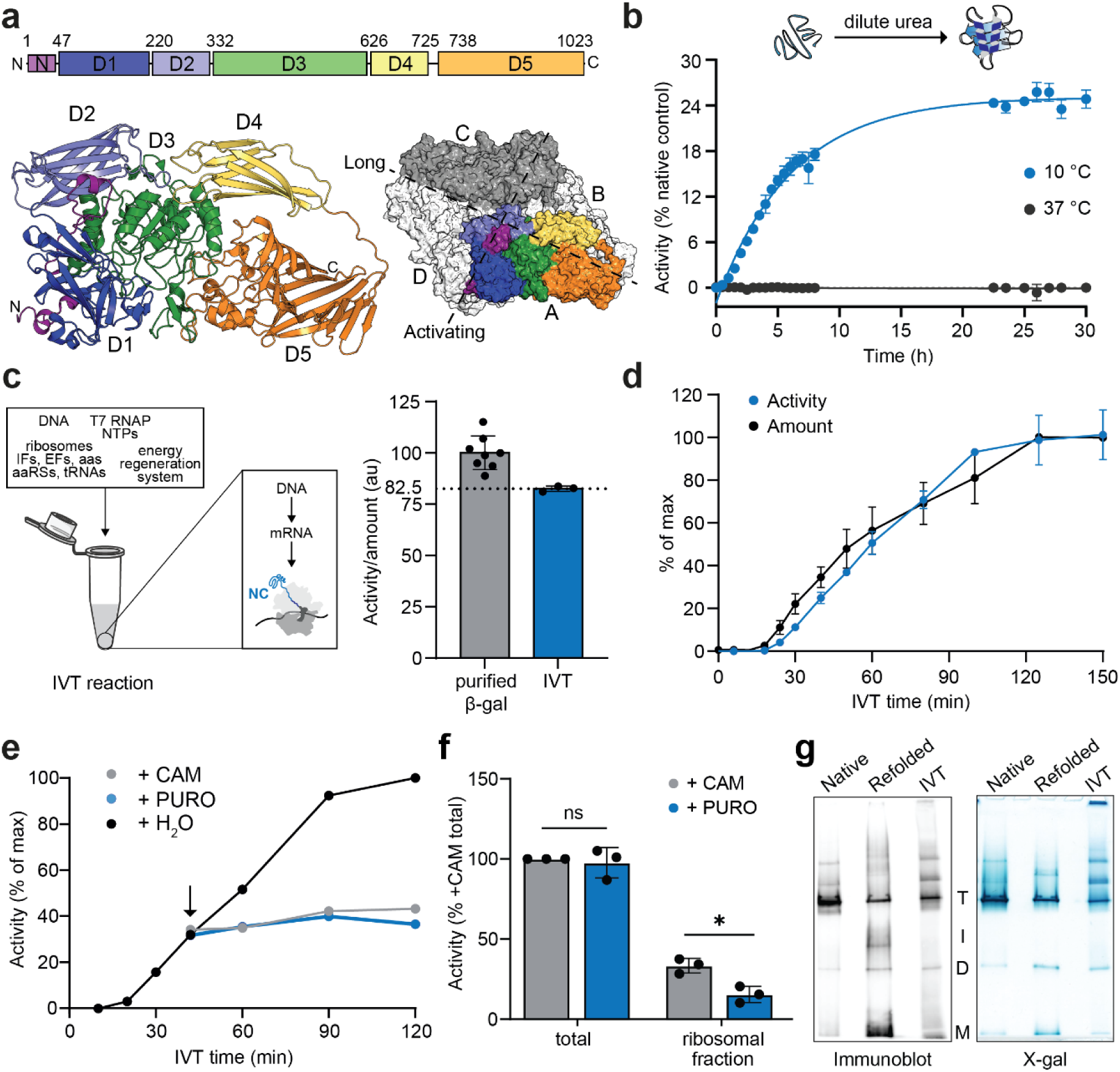
Cotranslational folding enables efficient biogenesis of β-gal. a, β-gal structure. Top, organisation of domains 1-5 (D1-5). Left, monomer structure (PDB: 6CVM^85^). Right, tetramer structure showing long and activating interfaces. **b,** Refolding of β-gal upon dilution from 8 M urea into buffer at 10 or 37 °C, assayed by recovery of enzyme activity. Data were fit to a single exponential function. Yield = 25.1%; 95% CI 24.5-25.7%. Half-time = 4.3 h; 95% CI 3.9-4.6 h. Error bars represent s.d. (n = 3). **c,** Specific activity of β-gal purified from *E. coli* (n=8 technical replicates) or produced by in vitro translation (IVT) (n=3 independent IVT reactions). Data were normalised to the mean specific activity of purified β-gal. Error bars represent s.d. **d,** Amount and activity of in vitro translated full-length β-gal, at different times after initiating IVTs using a plasmid encoding β-gal. Error bars represent s.d. (n = 3 independent IVT reactions). **e,** Accumulation of β-gal activity during IVT reaction. After 40 min (arrow), reactions were treated with 5 mM chloramphenicol (CAM), 5 mM puromycin (PURO), or an equal volume of water (H_2_O). **f,** β-gal activity in ribosomal fraction of IVTs. β-gal was expressed for 1 h before supplementing the reaction with either CAM to stabilise ribosome-nascent chain complexes, or PURO to release nascent chains. The ribosomal fraction was isolated by sucrose cushion ultracentrifugation. Error bars represent s.d. (n = 3 independent IVT reactions). ns - p>0.05; * - p<0.05 based on one-way ANOVA with Dunnett’s multiple comparisons. **g,** Oligomeric state of natively folded (native), refolded, or IVT-produced β-gal was analysed using native-PAGE. Duplicate gels were immunoblotted for β-gal or stained with the β-gal substrate X-gal. Bands corresponding to the tetramer (T), dimer (D), monomer (M) or intermediate (I) are indicated.

## RESULTS

### Cotranslational folding enables efficient biogenesis of β-gal

Spontaneous refolding of urea-denatured β-gal in vitro was undetectable at 37 °C, and inefficient (yield <30%) even at lower temperatures (Fig. 1b and S1a). Refolding was also slow, with t½ ∼4 h at 10 °C, and ∼10 min at 37 °C based on extrapolation of the Arrhenius plot (Fig. S1a,b). The bacterial Hsp70 chaperone system (DnaK, DnaJ, GrpE) supported only ∼20% refolding at 37 °C and did not increase the refolding rate (Fig S1c). β-gal refolding is therefore disfavoured at physiological temperature and only partially rescued by chaperones. In contrast, β-gal folded efficiently when produced in a fully-reconstituted in vitro transcription/translation (IVT) system without chaperones at 37 °C, to a yield of 83±1% active protein (Fig 1c). Moreover, several observations indicated that *de novo* folding of β-gal is rate-limited by its synthesis on the ribosome. First, the increase in β-gal enzyme activity during ongoing translation was near-concomitant with synthesis of the full-length protein (Fig 1d), consistent with previous work^28^. Second, β-gal activity did not increase after arresting translation using antibiotics (Fig 1e). Third, β-gal activity was detected in the ribosomal fraction of IVT reactions, and this was reduced by addition of puromycin to release nascent chains (Fig 1f). Thus, active β-gal accumulates on translating ribosomes, in agreement with previous in vivo observations^29–32^. Together, these data show that cotranslational folding of β-gal, even in the absence of chaperones, is both faster and substantially more efficient than its refolding from denaturant. We next sought to understand the molecular basis for this difference.

### Pathway of β-gal refolding from denaturant

We first aimed to understand why β-gal refolds to low yield *in vitro.* The absence of turbidity during refolding suggested that β-gal did not form large aggregates under our experimental conditions, although aggregation could be induced by raising the temperature and lowering the concentration of residual urea (Fig. S1d). To account for the fraction of β-gal that failed to refold, we resolved refolding reactions by native PAGE (Fig. S1e). Consistent with previous reports^33,34^, tetrameric β-gal accumulated over time at the expense of the monomer and dimer. We also observed a population of β-gal migrating as a diffuse band between dimer and tetramer on native PAGE, which appeared early during refolding and persisted throughout.

Prolonged incubation with X-gal revealed enzymatic activity of the monomer and dimer, indicating that they can assemble into active tetramers. However, the intermediate species was inactive (Fig. 1g and S1e). Thus, a fraction of β-gal becomes trapped as a non-native oligomer during attempted refolding, limiting the yield of native enzyme. In contrast, IVT-produced β-gal did not contain inactive intermediates (Fig. 1g). Cotranslational folding therefore bypasses a misfolding step that leads to off-pathway oligomerisation.

The fraction of β-gal that does refold from denaturant, refolds slowly. To understand why, we analysed refolding reactions using hydrogen/deuterium exchange-mass spectrometry (HDX-MS) (Fig. 2a). At 16 different times after dilution from denaturant, we pulse-labelled β-gal with D_2_O and quantified deuteron incorporation at the peptide level (Fig 2b,c). Our HDX-MS experiment followed the on-pathway refolding reaction, since the non-native oligomeric species was depleted during the deuterium labelling step (Fig S2a). We measured refolding kinetics for 400 individual peptides covering 96% of β-gal residues (Fig 2d, S2b and Data S1), allowing us to determine the detailed sequence of refolding events. The data were highly redundant, with each residue reported on by 4.4 peptides on average.

**Fig. 2.**
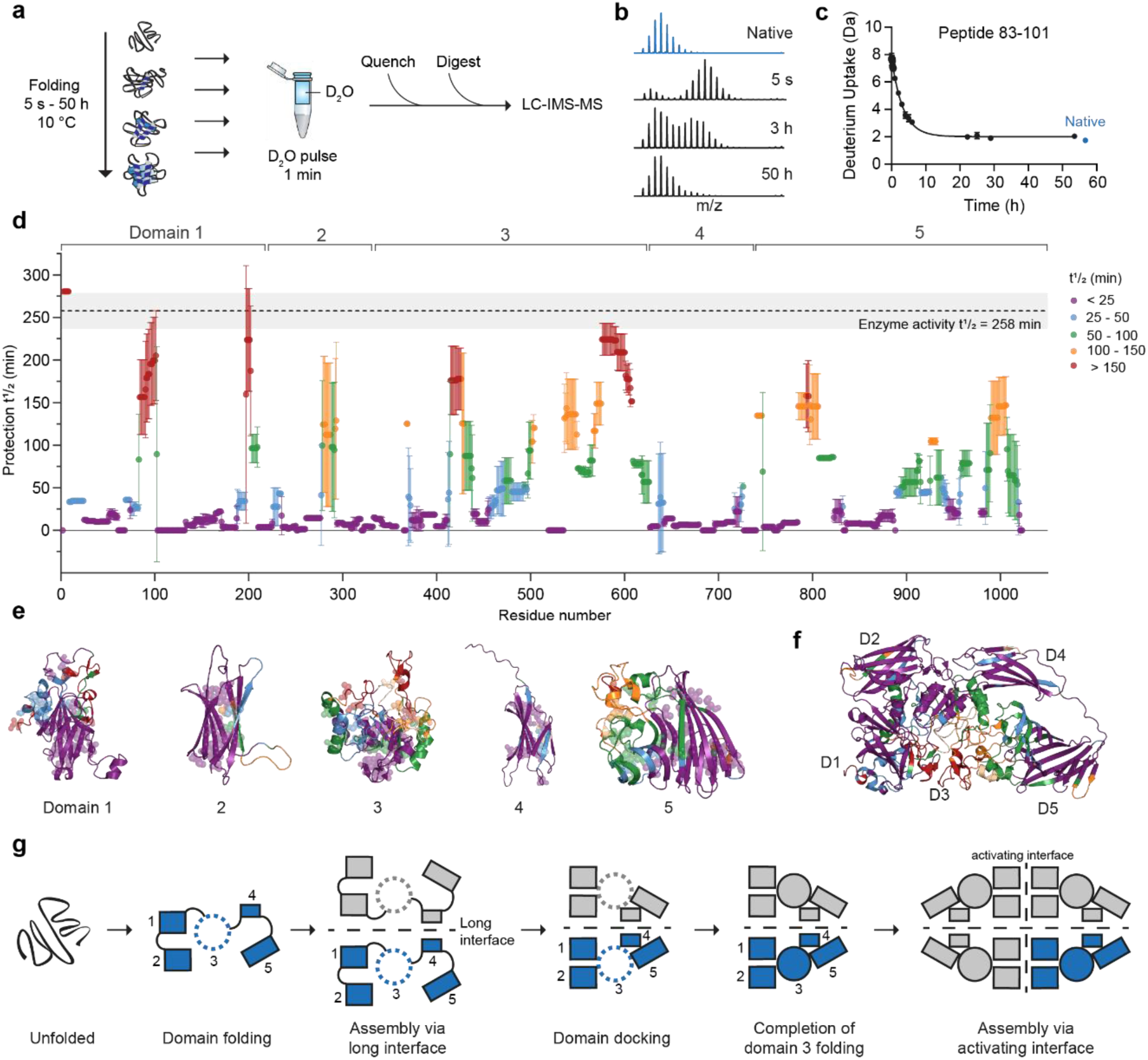
Pathway of β-gal refolding from denaturant. a, Scheme of HDX-MS experiment. **b,** Representative mass spectra for refolding of peptide 83-101, showing the time-dependent accumulation of the folded (protected) state at the expense of the unfolded (deprotected) state. **c,** Kinetics of peptide 83-101 refolding, calculated from the change in the centre of the spectral mass over time. Data were fit to a single exponential function. Error bars represent s.d. (n = 3). **d,** Half-times of HDX protection for peptides along the sequence of β-gal during refolding. The half-time and 95% CI for recovery of enzyme activity under the same conditions (Fig. 1b) are indicated. Error bars represent s.d. (n = 3). **e,** β-gal domains coloured by refolding half-time as in (**d**). Ile-Leu-Val clusters are shown as semi-transparent spheres. **f,** β-gal subunit coloured by refolding half-time as in (**d**). **g,** Schematic of β-gal refolding pathway.

β-sheet-rich regions in each domain, coinciding with hydrophobic Ile-Leu-Val clusters^35^, refolded early and near-simultaneously, followed by peripheral helices (Fig 2e). Slow refolding peptides tended to cluster in the native structure of β-gal, specifically at domain-domain interfaces involving D3 (Fig 2f and S3a,b). The long subunit interface reached native levels of exchange before domain interfaces within the monomer (Fig S3d,e), implying that dimer assembly initiates before monomer folding completes. This is consistent with previous analyses of β-gal refolding kinetics, which showed that the rate-limiting step is unimolecular and involves maturation of the dimer into a tetramerisation-competent state^33^. The slowest event was refolding of residues 3-8, comprising part of the α-peptide at the activating interface, which matches the kinetics of recovery of enzyme activity (Fig 2d and S3c). In summary, slow refolding of β-gal occurs in the following steps (Fig 2g): 1. Folding of the β-sheet core of each domain. 2. Assembly via the long interface. 3. Formation of domain:domain interfaces. 4. Completion of D3 folding. 5. Assembly via the activating interface. Non-native oligomerisation is a competitive side-reaction, possibly originating from partially-folded dimers that accumulate early during refolding.

### The ribosome directs segmental folding of the catalytic domain

We next sought to understand how the β-gal monomer folds on the ribosome. To model cotranslational folding intermediates, we prepared a series of 12 stalled ribosome:nascent chain complexes (RNCs), each representing an equilibrium snapshot of biosynthesis^36^ (Fig 3a,b and S4a,b). Translation was stalled at each domain boundary as well as within individual domains. To remove bound chaperones, we rigorously purified the RNCs from *E. coli* lacking the ribosome-associated chaperone Trigger factor. The three longest RNCs were also purified from Δ*lacZ* cells to remove low levels of co-purifying endogenous β-gal (Fig S4b and discussed below).

**Fig. 3.**
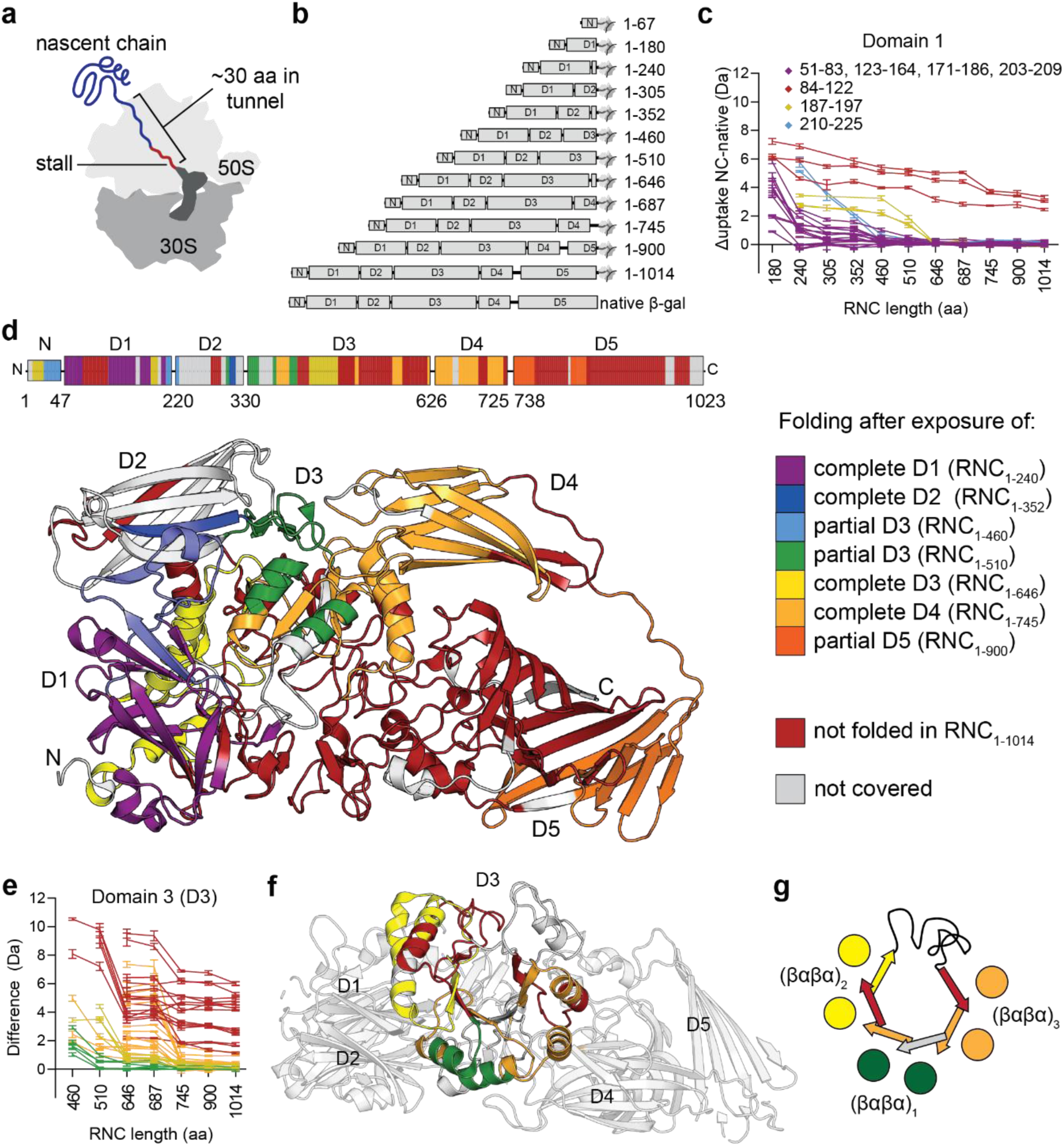
The ribosome directs segmental folding of the β-gal catalytic domain. a, Schematic of a stalled ribosome-nascent chain complex (RNC). Synthesis of the stalling sequence (WWWPRIRGPP, red) halts translation and allows isolation of stable RNCs of defined NC length. **b,** Schematic of β-gal RNCs, indicating NC length and domain boundaries. **c,** NC length-dependent protection of domain 1 peptides from deuterium exchange. The plot shows the difference in deuterium uptake, after 10 s deuteration, between peptides in indicated RNCs and full-length native β-gal. Peptides are grouped according to the NC length at which their level of deuterium uptake plateaus (i.e. changes <1 Da between successive RNCs). Error bars represent s.d. (n = 3). **d,** Cotranslational foldons. Peptides which plateau at the same NC length are grouped as in (**b**) and mapped onto the domain schematic (top) and monomer structure (bottom) of β-gal. Peptides that plateau without reaching the same level of uptake as native β-gal, even in the longest RNC, are coloured red and designated as “not folded” in RNC_1-1014_. Regions coloured white are not covered by any peptides. **e,** NC length-dependent protection of domain 3 peptides, as in (**b**). Peptide categories are mapped onto the structure of the β-gal monomer (**f**) and shown schematically (**g**).

The analytical complexity of ∼2.5 MDa RNCs poses a substantial challenge to probing NC conformation. We used an optimised HDX-MS workflow^8,36^ to define the folding status of each NC at the peptide level, using native β-gal as a reference (Fig S5 and Data S2). To uncover regions that fold cooperatively during synthesis, we plotted the uptake of each peptide as a function of NC length and grouped peptides that undergo their final folding transition at the same stage of synthesis (Fig 3c and S6a). We found that peptides grouped by folding behaviour also clustered spatially in the native structure of β-gal, indicating that they behave as cotranslational foldons (Fig 3d). Similar to the refolding reaction, β-rich regions often folded cooperatively on the ribosome. The β-sheet cores of D1, D2 and D4 reached near-native levels of deuterium exchange as soon as each complete domain was synthesised, as did a discontinuous β-rich subdomain in D5. In contrast, folding of loop- and α-helix-rich regions was often delayed relative to synthesis, suggesting a requirement for long-range stabilising interactions. For example, the central jelly roll of D1 folded as soon as the domain emerged from the ribosome, while peripheral loops and helices folded upon synthesis of neighbouring D2 and D3 (Fig 3c,d).

Cotranslational folding of D3 was distinct from other domains (Fig 3e-g and S6b). D3 houses the active site, and forms a distorted TIM-barrel with 3 βαβα repeats. Repeat 1 was synthesised in RNC_1-460_ but partially folded only in RNC_1-510_. Repeat 2 was synthesised in RNC_1-510_ and partially folded when D3 was complete in RNC_1-646_. Repeat 3 was synthesised in RNC_1-646_ and partially folded when D4 had emerged from the ribosome in RNC_1-745_. Thus, each βαβα repeat begins folding independently, following a short delay relative to their synthesis. Furthermore, the α-helices of each βαβα unit folded at shorter NC lengths than the β-strands, indicating that they may fold first during translation. This is in contrast to the refolding reaction, where the core β-strands of D3 mature near-simultaneously, followed by peripheral helices (Fig 2e). Together, these data define the sequence of cooperative folding events during β-gal synthesis, and reveal segmental, helix-driven folding of the D3 TIM-barrel.

### The ribosome surface sequesters a nascent amphipathic helix to alter domain folding

The central catalytic domain D3 refolds inefficiently (Fig 2), and populates unique intermediates during efficient folding on the ribosome (Fig 3). To understand how the ribosome modulates folding of this domain, we purified fragments containing the first 2¼ or 2½ domains of β-gal off the ribosome. HDX-MS showed that the 2¼-domain fragment was nearly identical to the corresponding RNC_1-460_, but the 2½-domain NC (RNC_1-510_) was conformationally destabilised on the ribosome as reflected by increased deuterium uptake (Fig S7a,b). Destabilised sites were distant from each other in sequence, but would cluster together near the C-terminus if the NC assumed a near-native tertiary structure (Fig S7c).

To understand the origin of ribosome-induced NC destabilisation, we determined the structure of RNC_1-510_ using cryogenic electron microscopy (cryo-EM) (Fig 4a and S8a-d). The overall architecture, with 70S ribosome bound to A- and P-site tRNAs, is similar to previously reported structures of stalled ribosomes^37^. In addition, we observed continuous density for nascent β-gal in the tunnel and against the ribosome surface, of sufficient quality to model both the stalling sequence and the C-terminal 46 residues (465-510) of the β-gal construct (Fig. 4b). Residues 485-494 near the vestibule were modelled as an alanine backbone due to limited local resolution. The NC assumes an extended conformation in the exit tunnel and follows a defined path, stabilised by interactions with ribosomal proteins uL4 and uL22 lining the tunnel wall (Fig. S9a-d). Although it initially follows the contour of uL22, the NC switches trajectory near the vestibule, towards uL29, before emerging from the tunnel close to uL23 (Fig. 4a and S9b). Density corresponding to the N-terminal part of the NC (residues 1-464) is visible only at lower contour levels (Fig 4a).

**Fig. 4.**
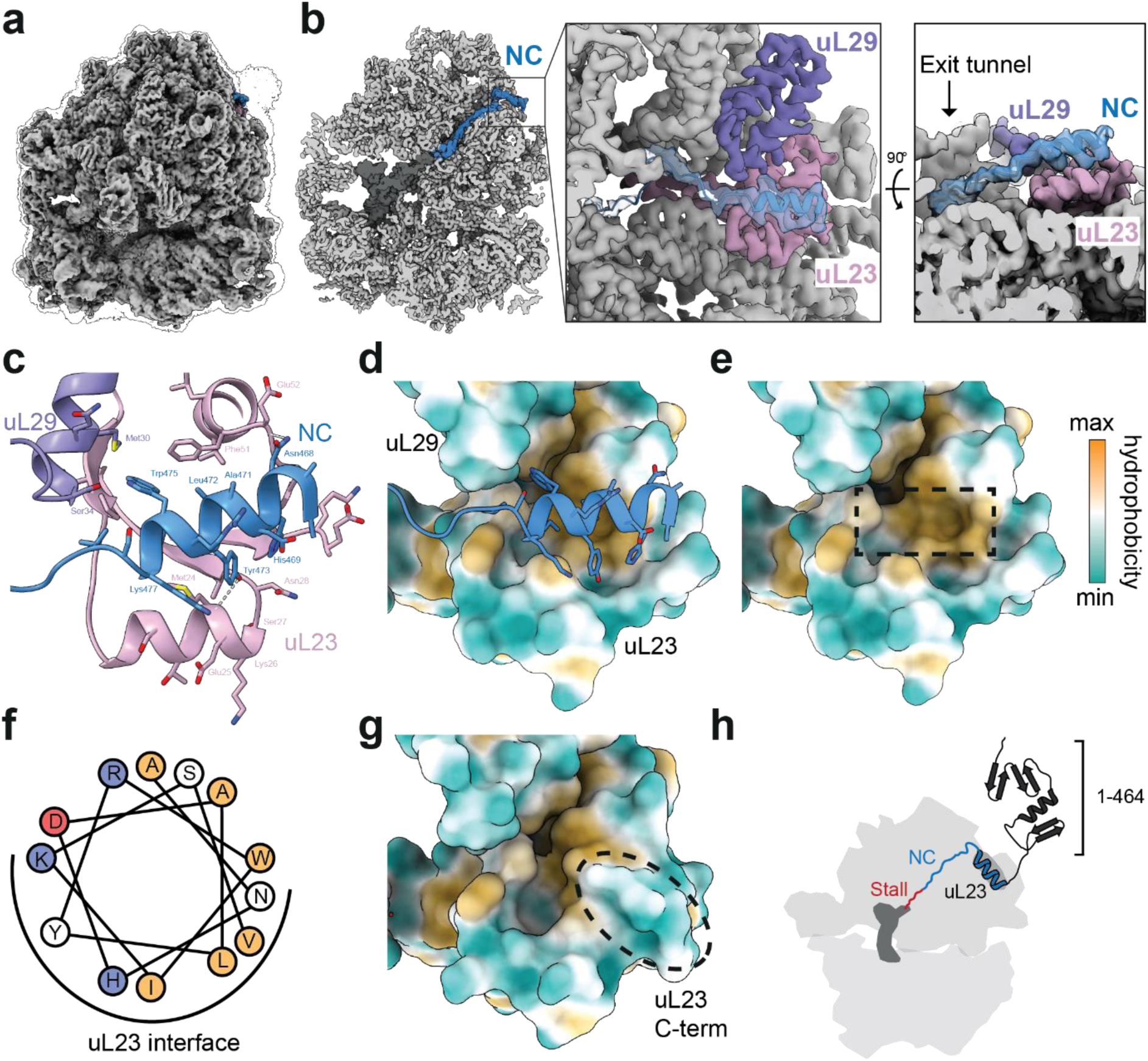
The ribosome surface sequesters a nascent amphipathic helix to alter domain folding. a, Cryo-EM structure of RNC_1-510_. The silhouette outlines the same map at a lower contour level. **b,** Cross-section of the map, with the peptidyl-tRNA coloured dark grey and the nascent chain (NC) coloured blue. Inset: close-up view of NC interactions with uL23 (salmon) and uL29 (purple). NC density is semi-transparent and the NC model is shown in cartoon representation. **c**, Interactions between the NC and uL23/uL29. **d**, Model of NC (blue) bound to the ribosome surface, with uL23 and uL29 surfaces coloured by hydrophobicity. **e**, as in (**d**), with the NC removed. The position of the NC helix is indicated by a dashed box. **f**, Helical wheel representation for residues _466_ANHDALYRWIKSV_478_, coloured according to electrostatic potential and hydrophobicity (red - negative, blue - positive, hydrophobic - orange). **g**, Surface hydrophobicity of uL23 and uL29 in the 70S ribosome without a nascent chain (7K00^72^). The C-terminus of uL23 (residues 89-100) is indicated with a dashed box. **h**, Model for ribosome-induced destabilisation of RNC_1-510_. The ribosome sequesters residues 466-478 via a hydrophobic groove on uL23, preventing this region from forming native contacts with the N-terminal part of the NC.

Strikingly, residues 466-478 of β-gal fold into a well-defined α-helix (local resolution 2.8-3.0 Å) outside the vestibule (Fig 4b,c and S8e). The helix is anchored by NC residues Tyr473 and Asn468, which form hydrogen bonds to uL23 residues Met24 and Glu52, respectively. Trp475 of the NC inserts into a pocket formed by Ser34 and Met30 of uL29, and Phe51 of uL23. The groove formed by uL23/uL29 is hydrophobic and accommodates the nonpolar face of the amphipathic NC helix (Fig. 4d-f). Although residues 466-478 are α-helical in native β-gal, we found that the equivalent isolated peptide is disordered (Fig S9e,f), suggesting that helical structure is stabilised by uL23. Interestingly, the NC-binding pocket we identify is normally occupied by the flexible C-terminus of uL23, which is unresolved in our structure and presumably displaced by the NC (Fig. 4g). Residues that line the binding pocket and confer its hydrophobic character are evolutionarily conserved (Fig. S9g).

Folding of the nascent helix in complex with uL23 explains why NC_1-510_ is locally destabilised on the ribosome. In native β-gal, the hydrophobic face of the helix packs against a hydrophobic groove in D3 (Fig S9h). Interactions with the ribosome therefore compete with intra-NC interactions, disrupting NC folding (Fig. 4h). Since the destabilised sites in RNC_1-510_ form part of the same cotranslational foldon (Fig. 3c and S7b,c), helix sequestration suffices to destabilise adjacent regions. Importantly, the locally destabilised regions recover when the chain elongates further in RNC_1-646_, indicating that intra-NC interactions eventually outcompete ribosome binding (Fig. S7b). In summary, we show that the ribosome surface can modulate domain folding by temporarily sequestering a nascent amphipathic helix in a cryptic hydrophobic pocket. This results in segmental cotranslational maturation of a domain that otherwise refolds in a concerted fashion.

### The ribosome directs β-gal assembly to avoid misassembly

During refolding, β-gal assembles prematurely and forms off-pathway oligomers (Fig 1 and 2). In contrast, assembly is efficient when coupled to ongoing translation^38^ (Fig. 1c-g). We therefore considered whether the ribosome changes the pathway of β-gal assembly to promote productive folding. Since β-gal is only active when tetrameric, the observation of β-gal activity on polysomes implies that it assembles to some extent during synthesis (Fig 1f). To probe the underlying mechanism, we first measured enzyme activity of purified β-gal RNCs (Fig 5a). RNCs longer than 745 aa showed low levels of enzyme activity (<5% of RNC_FL+50aa_), indicative of weak or partial assembly. Assembly involved endogenous β-gal, since RNCs produced in Δ*lacZ* cells or IVTs were inactive (Fig 5b and S10a).

**Fig. 5.**
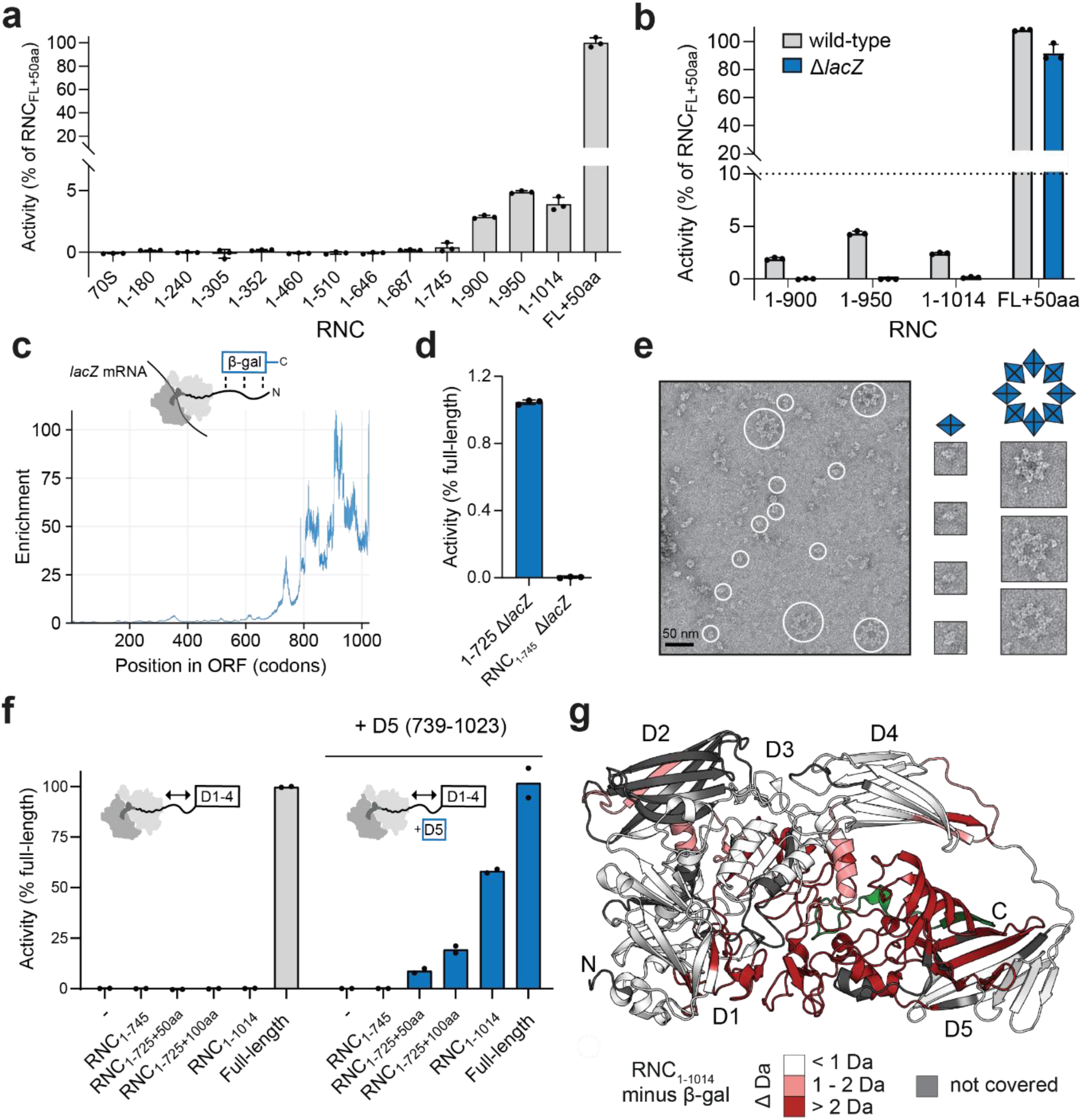
The ribosome directs β-gal assembly to avoid misassembly. a, β-gal enzyme activity of 5 nM RNCs purified from wild-type *E. coli*. Error bars represent s.d. (n = 3). **b,** Endogenous β-gal assembles with RNCs. Enzyme activity of 5 nM RNCs purified from wild-type, or Δ*lacZ E. coli* which do not express β-gal. Error bars represent s.d. (n = 3). **c,** Onset of co-post β-gal assembly in vivo. SeRP data showing the codon-resolved enrichment (selected translatome / total translatome) of β-gal-bound RNCs along the *lacZ* open reading frame. **d,** D1-4 are active off but not on the ribosome. Enzyme activity of 120 nM 1-725 β-gal truncation (domains 1-4) and RNC_1-745_, both purified from Δ*lacZ E. coli*. Error bars represent s.d. (n = 3). **e,** Truncated β-gal misassembles. Negative-stain EM micrograph of β-gal 1-725 purified from Δ*lacZ E. coli.* Particles corresponding to tetramers or higher-order oligomers are indicated with blue or white circles, respectively. **f,** β-gal assembly is triggered by folded D5. Enzyme activity of 7.5 nM β-gal RNCs purified from Δ*lacZ E. coli*, in the absence or presence of isolated folded domain 5 (D5, residues 739-1023). The distance between D1-4 (residues 1-725) and the ribosome was increased by inserting a 50 or 100 aa Gly-Ser linker between the C-terminus of the NC and the stalling sequence. **g,** D5 folds primarily post-translation. Difference in deuterium uptake, after 100 s deuteration, between native β-gal and RNC_1-1014_ purified from Δ*lacZ E. coli*. Regions that are deprotected in the RNC are coloured in shades of red. The C-terminal 30 aa, expected to be inside the exit tunnel in RNC_1-1014_, are coloured green.

Different assembly modes are possible, depending on whether assembly initiates between two incomplete subunits (“co-co” translational) or one incomplete and one complete subunit (“co-post” translational)^21^. Consistent with co-post assembly, RNCs longer than 745 aa co-purified with full-length β-gal (Fig S4a). This occurred in WT but not Δ*lacZ* cells, and was partially disrupted by mutating the activating interface (Fig S4b, and S10b). The co-assembling full-length subunits were protease resistant, suggesting that they are folded (Fig S10c). Thus, mature β-gal subunits assemble with incomplete nascent polypeptides exposing at minimum the first 4 domains of nascent β-gal.

To test this mechanism in vivo, we used selective ribosome profiling^21^. Pull-downs using C-terminally tagged β-gal efficiently recovered ribosomes translating *lacZ* mRNA, indicative of co-post assembly (Fig. S10d). Analysis of protected footprints revealed that the onset of assembly occurred after synthesis of ∼700 aa, stabilised after ∼800aa, and persisted until translation termination, in close agreement with the pulldowns of stalled RNCs (Fig. 5c). The late onset of assembly implied that co-post assembly initiates via the long (C-terminal) rather than activating (N-terminal) interface. Consistent with this, a β-gal fragment 1-519, which contains all residues contributing to the activating interface, was monomeric even at a concentration of 900 µM (Fig S10e).

RNCs were inactive when produced in vitro in the absence of full-length subunits, arguing against a co-co assembly pathway involving two or more incomplete NCs (Fig S10a). To detect possible co-co assembly *in vivo*, we performed disome selective profiling^20^ (Fig S10f,g). Co-co assembly is expected to shift translating ribosomes from the monosome to disome fraction after nuclease digestion of the connecting mRNA, resulting in an enrichment of disome footprints at the expense of monosome footprints. No such enrichment was observed for β-gal, indicating that cotranslational β-gal assembly is co-post.

To test whether incomplete chains of β-gal can self-assemble off the ribosome, we purified a 4-domain fragment (1-725), corresponding to the earliest onset of assembly detected by SeRP. We found that 1-725 formed unstable tetramers with weak enzymatic activity (Fig 5d and S10h). However, negative-stain EM revealed additional higher-order oligomers containing 7-8 copies of 1-725 tetramers arranged in a star-like structure (Fig 5e and S10i). Incomplete chains of β-gal are therefore assembly-competent off the ribosome, but can misassemble. In contrast to isolated 1-725, RNC_1-745_ (purified from Δ*lacZ* cells) was inactive and thus unassembled (Fig 5d). The ribosome therefore inhibits premature co-co assembly, and in doing so prevents misassembly.

To test whether the ribosome sterically interferes with co-co assembly, we purified RNCs exposing residues 1-725 of β-gal fused to C-terminal Gly-Ser linkers of increasing length. These RNCs were inactive, but could be activated in a linker length-dependent manner by complementation with folded D5 *in trans* (Fig 5f). Cotranslational assembly is therefore influenced by both subunit folding and proximity to the ribosome.

To understand how folding and assembly are coordinated prior to translation termination, we focused on RNC_1-1014_. This RNC represents the endpoint of cotranslational maturation, when only ∼30 C-terminal residues remain sequestered in the ribosome exit tunnel. HDX-MS analysis of RNC_1-1014_ showed that D5 was strongly destabilised compared to native β-gal, and must therefore complete folding post translation (Fig 5g and S5). Since D5 forms the majority of the long interface, post-translational D5 folding may explain why cotranslational assembly strictly requires at least one full-length subunit. Indeed, RNC_1-1014_ was efficiently activated by purified D5 in *trans*, demonstrating that D5 folding can trigger co-co tetramer assembly on ribosomes (Fig 5f). The ribosome therefore dictates the mode and onset of assembly by controlling the timing of C-terminal domain folding. Assembly is synchronised with monomer folding, rather than preceding folding as occurs during inefficient refolding from denaturant (Fig 2 and S3).

## DISCUSSION

By characterising the maturation of a large multidomain oligomeric protein on and off the ribosome, we uncover new features of cotranslational folding that support the biogenesis of complex folds. We show that the ribosome directs segmental folding of a nascent TIM barrel, and uses a conserved hydrophobic groove on L23 to change the order of folding events such that α-helices fold first (Fig. 6). Since isolated secondary structures are intrinsically unstable, nucleating helix folding may promote folding by reducing the entropic barrier^5,39,40^. Indeed, folding of a model protein could be accelerated by stabilising native-like helical contacts^41^. Our results demonstrate that the ribosome surface is not necessarily generically destabilising^9,11,42^, but can direct specific folding outcomes. Although it is not yet clear whether this phenomenon is general, we note that nascent apomyoglobin was found to dock close to the same hydrophobic pocket on uL23^43^.

**Fig. 6.**
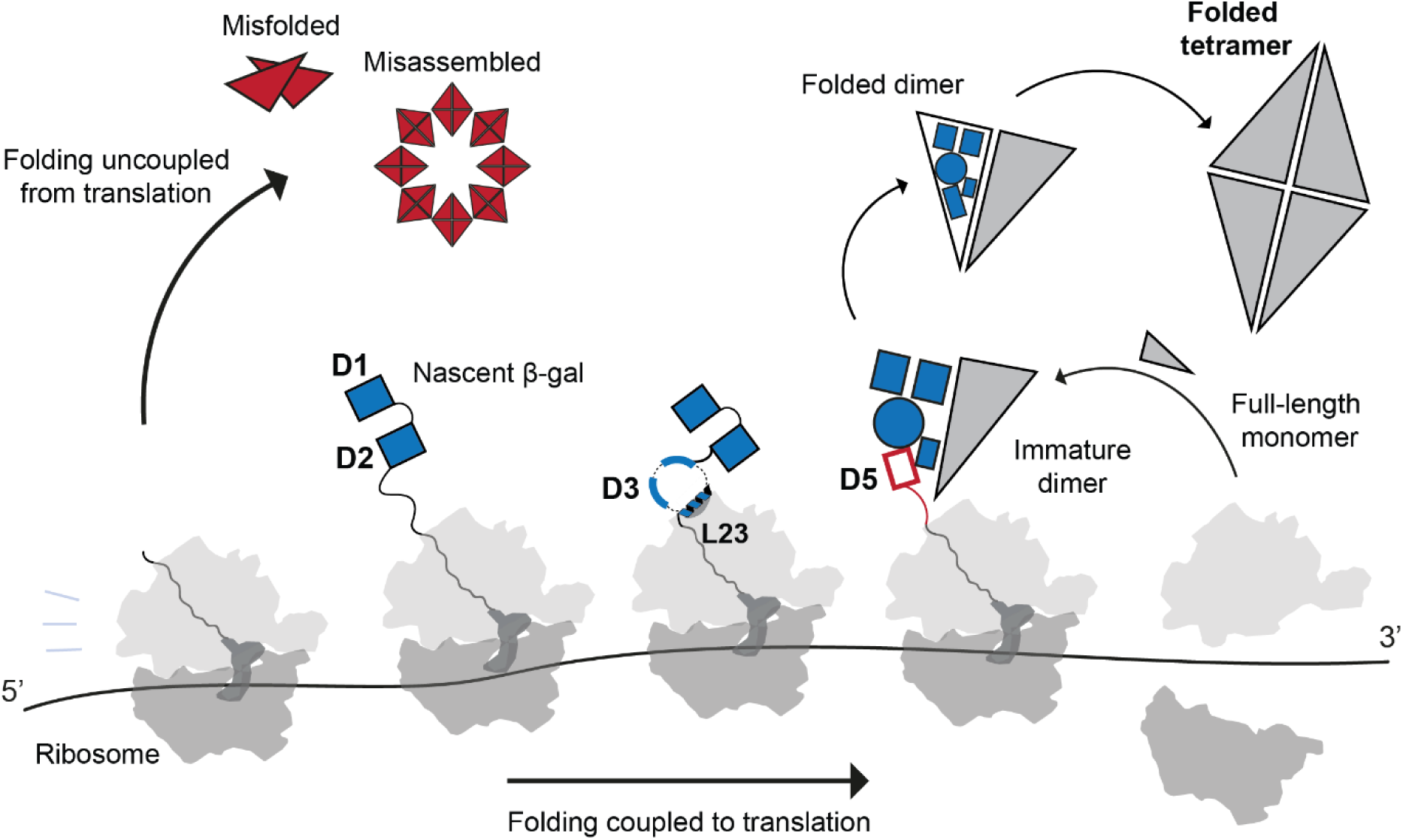
Cotranslational folding and assembly of β-gal. β-gal folds efficiently when coupled to ongoing translation, directed by the ribosome to synchronise with tetramer assembly. Domain 3 folds in discrete steps, the timing of which are dictated by an interaction with a hydrophobic groove on uL23 that stabilises a nascent amphipathic helix. The pioneering round of translation generates a full-length subunit, which then binds via the long interface to a ribosome-associated subunit during synthesis of domain 5. An immature dimer results, which matures post-translation and assembles via the activating interface to form the folded tetramer. When uncoupled from translation, monomers fold via a less efficient pathway and are prone to misfold and misassemble.

Approximately 10% of proteins contain TIM barrels^44^ and this domain is enriched among substrates of eukaryotic and bacterial chaperonins^45,46^. Interestingly, the segmental, helix-first folding we observe on the ribosome is also characteristic of accelerated TIM barrel folding in the bacterial chaperonin GroEL/ES^47^. Furthermore, GroEL has been shown to stabilise α-helical conformations of bound peptides^48,49^. uL23-directed folding may constitute an alternative, chaperone-independent mechanism supporting TIM barrel biogenesis.

Oligomers present a special biogenesis challenge, as assembly must coordinate with folding while avoiding spurious misassembly^24,50–52^. Nonetheless, most proteins across all proteomes are oligomeric^3^, implying that oligomer biogenesis is optimised in vivo. We find that the ribosome specifies the pathway of β-gal assembly. Although β-gal is a symmetric homomer, cotranslational assembly is directional and involves a full-length and a partially-synthesised subunit (Fig. 6). This assembly mode may help to confer assembly specificity when multiple oligomeric forms are possible^53,54^ (Fig 5e), and avoid misfolding between two incompletely folded subunits^50^ (Fig 1g). Since β-gal tetramers are extremely stable^55,56^, the co-assembling full-length subunit is unlikely to originate from mature tetramers in vivo. Instead, we propose that the pioneering round of translation generates a subunit which then engages the nascent chain on the trailing ribosome (Fig. 6). The polysome may therefore act as a scaffold for efficient co-post homomer assembly. Notably, co-co assembly occurs for only a subset of homomers, characterised by intertwined subunits^18,20,57^. Future work will need to establish the prevalence of the alternative co-post homomer assembly mode we describe here.

In summary, we show how the translating ribosome provides a platform for biogenesis of proteins with complex topologies. Together with the chaperone network, this may have shaped the evolution of protein folds by allowing domains and subunits to be combined without compromising foldability.

## ACKNOWLEDGEMENTS

We thank the Francis Crick institute Proteomics STP (Sarah Maslen and Mark Skehel for help with HDX-MS), the Chemical Biology STP (Dhira Joshi for peptide synthesis, Christelle Soudy for help preparing pepsin beads), the Structural Biology STP (Andrea Nans for cryo-EM data collection, Qu Chen for nsEM training, Laura Masino for help with CD), the Scientific Computing STP for support with high performance computing, Milos Cvetkovic for help with nsEM grid preparation, Stephane Mouilleron for help with SEC-MALLS, and all members of the Protein Biogenesis and Visual Biochemistry laboratories for help and discussion. B.B. acknowledges a research grant from the European Union (ERC - SyG - 101072047 - CoTransComplex). Views and opinions expressed are however those of the authors only and do not necessarily reflect those of the European Union or the European Research Council. Neither the European Union nor the granting authority can be held responsible for them. J.S. was supported by fellowships from the Studienstiftung (German Academic Scholarship Foundation) and the Baden-Württemberg State Graduate Funding (Landesgraduiertenförderung, LGF). J.S. is a member of the Heidelberg Biosciences International Graduate School (HBIGS). D.B.’s work is supported by the Francis Crick Institute which receives its core funding from Cancer Research UK (CC2025), the UK Medical Research Council (CC2025), and the Wellcome Trust (CC2025), and by UK Research and Innovation (UKRI) under the UK government’s Horizon Europe funding guarantee [FoldingMap, EP/X020428/1]. Work in the Enchev laboratory is supported by the Francis Crick Institute, with funding from Cancer Research UK (CC2059), the UK Medical Research Council (CC2059), and the Wellcome Trust (CC2059) as well as a Royal Society Wolfson Fellowship and The Crick Chris Banton Translation Fund.

## AUTHOR CONTRIBUTIONS

A.R. performed the biochemical and HDX-MS analyses of RNCs. S.S. performed the biochemical and HDX-MS analyses of refolding reactions. G.J. and J-Z.E. collected and analysed cryoEM data with help from R,I.E. J.A., J.L.S., G.K. and B.B. performed and analysed the RiboSeq experiments. D.B. and R.I.E conceived and supervised the project, and wrote the manuscript together with A.R., S.S., G.J. and J-Z.H.

## SUPPLEMENTARY DATASETS

**Data S1**. HDX-MS analysis of β-gal refolding.

**Data S2**. HDX-MS analysis of β-gal truncations and RNCs.

**Data S3**. Sequences of recombinant proteins used in this study.

## METHODS

### DNA vectors

Full-length and truncated *E. coli* β-galactosidase variants were expressed from a pET28 vector following an N-terminal His_6_-tag and TEV-protease site^58^. RNCs were expressed from pET21 plasmids encoding β-galactosidase ORFs upstream of an arrest-enhanced ribosome stalling sequence (WWWPRIRGPP)^58,59^. *E. coli* DnaK and DnaJ were expressed without any tags from pET11d. *E. coli* GrpE was expressed without any tags from pET3a. Chaperone expression vectors were kind gifts from F.U. Hartl (MPI Biochemistry). Additional mutations were introduced using site-directed mutagenesis with Q5 polymerase (NEB, E0554). All constructs used in this study were verified by sequencing and are listed in Data S3.

### Expression and purification of full-length β-gal

*E. coli* BL21DE3 cells were transformed with a pET28 vector encoding β-gal fused to a N-terminal His_6_-tag followed by a TEV protease recognition site. An overnight bacterial culture was used to inoculate Terrific broth, and the cells were grown at 37 °C to an OD of 0.8. β-gal expression was induced with 1 mM isopropyl β-D-1-thiogalactopyranoside (IPTG) and further incubation was at 16 °C for 18 h. The cells were harvested by centrifugation and resuspended in 50 mM Tris-HCl pH 7.5, 500 mM NaCl, 10 % (v/v) glycerol, 2 mM β-mercaptoethanol and 0.1 mM PMSF (Buffer A). Just before lysis, cells were supplemented with benzonase, 1 mM PMSF and lysed by two passes through a French press (Constant Systems) operating at maximum pressure of 25 kpsi and 4 °C. The clarified supernatant was loaded on to a 5 ml HisTrap HP (Cytiva) column using an AKTA pure system (Cytiva). The column washed with Buffer A, and the proteins were eluted using a linear gradient of 0-500 mM imidazole in Buffer A. Fractions with β-gal were pooled, supplemented with TEV protease, and dialyzed overnight against 20 mM Tris-HCl pH 7.5, 250 mM NaCl, 5 mM MgCl_2_, 10 % (v/v) glycerol and 2 mM DTT at 4 °C. β-gal that was cleaved off the His tag was separated by passing the dialyzed protein through a 5 ml HisTrap HP column. The protein was concentrated, and injected into a HiPrep 26/60 Sephacryl S-300 HR column (Cytiva) equilibrated with 20 mM Tris-HCl pH 7.5, 300 mM NaCl, 10 mM MgCl_2_, 5 % (v/v) glycerol and 3 mM DTT. Fractions containing pure protein were concentrated, flash frozen in liquid nitrogen and stored at -70 °C.

### RNC buffers

RNC low-salt buffer contained 50 mM HEPES-NaOH pH 7.5, 12 mM Mg(OAc)_2_, 100 mM KOAc, 1 mM DTT and 8 U/mL RiboLock RNase inhibitor (ThermoScientific, EO0384). RNC high-salt sucrose cushion contained 35% sucrose, 50 mM HEPES-NaOH pH 7.5, 12 mM Mg(OAc)_2_, 1 M KOAc, 1 mM DTT, 8 U/mL RiboLock RNase inhibitor and 0.2x Halt Protease Inhibitor Cocktail (ThermoScientific, 78437). RNC low-salt sucrose cushion contained 35% sucrose, 50 mM HEPES-NaOH pH 7.5, 12 mM Mg(OAc)_2_, 100 mM KOAc, 1 mM DTT, 8 U/mL RiboLock RNase inhibitor and 0.2x Halt Protease Inhibitor Cocktail.

### Expression and purification of ribosome:nascent chain complexes (RNCs)

RNCs were expressed and purified as described previously^58^ with slight modifications. In short, BL21(DE3) *E. coli* cells were transformed with plasmids encoding RNC constructs and grown in ZYM-5052 autoinduction media^60^ for 18 h at 37 °C. As indicated, expression cells were either TF knock-out cells (*Δtig,* John Christodoulou, UCL) or β-gal knock-out cells (*ΔlacZ*, Addgene, 99247^61^). Cells were pelleted (4,000 g, 30 min) and resuspended in RNC lysis buffer (70 mM Tris, pH 7.5, 150 mM KCl, 10 mM MgCl_2_, 8 U/mL RiboLock RNase inhibitor, 0.5x Halt Protease Inhibitor Cocktail, 2 mg/mL lysozyme, 0.05 Kunitz units/μL RNase-free DNase (QIAGEN, 79254)). Following an incubation (30 min, 4 °C) and subjecting resuspended pellets to at least 2 freeze-thaw cycles at -80 °C, the soluble fraction was separated by centrifugation (20 min, 16,000 g) and centrifuged (2 h, 264,000 g) through RNC high-salt sucrose cushion to isolate ribosomes in the pellet. The pellet was resuspended in RNC low-salt buffer and applied to in-house prepared GFP-clamp-agarose beads^62^ for 16 hours at 4 °C. RNCs were selectively eluted by muGFP-tag cleavage with HRV 3C protease, and further purified by pelleting through a second sucrose cushion (2 h, 264,000 g). Pellets were resuspended in RNC low-salt buffer, snap-frozen in liquid nitrogen and stored at −80 °C. Sequences of all purified RNCs are listed in Data S3.

### Expression and purification of β-galactosidase truncations

Expression and purification of β-galactosidase truncations was performed as described previously^58^ with slight modifications. BL21(DE3) *E. coli ΔlacZ* cells (Addgene, 99247^61^) transformed with plasmid encoding C-terminally truncated β-gal 1-440, 1-519, 1-725 or β-gal domain 5 (D5, 739-1023) were grown in LB at 37 °C until OD_600_ 0.6–0.8 and protein expression was subsequently induced with 1 mM IPTG for an additional 18 hours at 16-18 °C. BL21(DE3) *E. coli ΔlacZ* cells transformed with plasmid encoding C-terminally truncated β-gal 1-490 were grown in ZYM-5052 autoinduction media ^60^ for 18 h at 37 °C. Cells were harvested (4,000 g, 30 min), resuspended in Buffer A (50 mM Tris-HCl, pH 7.5, 500 mM NaCl, 10% glycerol, 2 mM β-mercaptoethanol, 100 mM PMSF) supplemented with 1 mg/mL lysozyme, benzonase (Millipore, E1014) and Complete EDTA-free protease inhibitor cocktail (Roche, 11873580001), and lysed by sonication. Lysate was clarified by centrifugation (60,000 g, 45 min) and then applied to HisTrap HP column (Cytiva, 17524802) equilibrated in Buffer A. Peak fractions from gradient elution with 500 mM Imidazole were treated with TEV protease (48 h, 4 °C) and dialysed against Buffer A. Uncleaved protein was removed using a HisTrap HP column equilibrated in Buffer A. The flow-through was further purified using a Superdex 200i 10/300 (Cytiva, 28990944) equilibrated in Buffer B (20 mM Tris-HCl, pH 7.5, 300 mM NaCl, 10 mM MgCl_2_, 3 mM DTT, 5% glycerol), and pure protein was concentrated and snap-frozen in liquid nitrogen for storage at −80 °C. Sequences of purified proteins are listed in Data S3.

### Expression and purification of DnaK, DnaJ and GrpE

DnaK, DnaJ and GrpE were expressed and purified as described previously^58^ without any modifications. Sequences of purified proteins are listed in Data S3.

### SEC-MALS

100 μL of pure protein was applied at 0.5 mL/min to Superose 6i 10/300 GL column (Cytiva) equilibrated in 30 mM Tris-HCl, pH 7.5, 10 mM MgCl_2_, 300 mM NaCl, 2 mM DTT for an isocratic elution in the same buffer. The concentrations of injected proteins are indicated in figure legends and were determined using absorbance at 280 nm and the Expasy ProtParam predicted extinction coefficients^63^. UV absorbance, light scattering (LS) and differential refractive index (dRI) were monitored throughout elution. The molecular mass of the eluate was determined from LS and dRI data using the ASTRA software.

### Immunoblotting

Following SDS-PAGE, proteins were transferred onto a PVDF membrane using a Trans-Blot Turbo Transfer System (BioRad). Membranes were blocked in PBS-Tween with 5% non-fat milk for 1 hour at RT and incubated with appropriate primary antibodies (1:1,000 dilution in PBS-Tween with 5% non-fat milk) for 1 hour at RT. Following three washes (5 min, RT) with PBS-Tween, membranes were incubated with appropriate HRP-conjugated secondary antibodies (1:10,000 dilution in PBS-Tween with 5% non-fat milk) for 1 hour at RT and washed. Membranes were developed by enhanced chemiluminescence using SuperSignal West Pico PLUS Chemiluminescent Substrate (ThermoScientific, 34080). Primary antibodies used for each immunoblot are specified in figure legends – these include monoclonal mouse anti-β-galactosidase antibody (Santa Cruz Biotechnology, antibody sc-65670, 1:1000) and polyclonal chicken anti-β-galactosidase N-terminus antibody (abcam, ab106567, 1:1000). These were detected with corresponding HRP-conjugated secondary antibodies – either goat polyclonal anti-mouse (abcam, ab205719) or anti-chicken (abcam, ab97135). Quantification of band-specific chemiluminescent signal was performed in Fiji^64^ by measuring the area under a signal peak corresponding to the given band.

### β-galactosidase refolding

β-gal refolding experiments were performed essentially as described previously^33^. The concentration of β-gal was estimated using the Bradfords assay reagent (Bio-Rad) with Bovine Serum Albumin Standard (ThermoFisher). The unfolding reaction containing 10 µM β-gal in 100 mM sodium phosphate pH 7.5, 8 M urea, 1 mM EDTA and 10 mM DTT was incubated overnight at 10 °C. For the native control, β-gal was incubated in the same buffer without urea. Refolding reactions were initiated by diluting the unfolding reaction (or native control) 10-fold into the refolding buffer containing 100 mM sodium phosphate pH 7.5, 0.7 M urea, 1 mM MgCl_2_ and 5 mM DTT at 10 °C. To measure the extent of refolding, 1 µl of the refolding reaction was added to 300 µl of reaction mix containing 30 mM TES pH 7.0, 145 mM NaCl, 10 mM MgSO_4_, 0.1 mM EDTA and 2 mM o-nitrophenyl-β-D-galactopyranoside. The increase in absorbance at 420 nm was measured using Spectramax Plus 384 plate reader (Molecular Devices) operating at 23 °C and the slope of the progress curve was recorded. For refolding in presence of Hsp70 system, the refolding as well as the native control reaction contained 100 mM Hepes-sodium pH 7.5, 10 mM MgCl_2_, 100 mM KCl, 5 mM DTT, 3 µM DnaK, 1 µM DnaJ, 1.5 µM GrpE and 2.5 mM ATP. Refolding yield (ratio of activity of refolded β-gal to the activity of the native control) was plotted against time in Prism (GraphPad Software), and the data were fit to a one-phase association model to determine the half-time and rate constant of refolding.

### β-galactosidase RNC activity assay

Enzymatic activity of β-gal-containing IVTs, β-gal RNCs and β-gal C-terminal truncations was measured in RNC low-salt buffer with final 2.1 mM o-nitrophenyl-β-D-galactopyranoside (oNPG) by following absorbance at 420 nm at 25 °C and recording the slope of the progress curve. See figure legends for experiment-specific information about final protein concentrations. Unless otherwise stated, reactions were diluted 200-fold in oNPG-containing buffer just prior to activity measurement.

### In vitro translation

IVT reactions were performed using the PURExpress *in vitro* Protein Synthesis Kit (NEB, E6800S). Solutions A and B were pre-mixed according to manufacturer’s instructions at 2:1.5 ratio on ice, and all reactions were supplemented with RNasin Ribonuclease Inhibitor (Promega, N2111) at a final concentration of 2 units/μL. Any supplemented plasmid DNA was added at final 10 μg/μL concentration. For continuous IVT monitoring, fluorogenic resorufin-β-D-glucopyranoside (Merck, R4883, λ_ex_ 570 nm, λ_em_ 585 nm) was added at final 250 μM. Unless otherwise stated, expression was subsequently conducted at 37 °C for up to 2 hours. Expressed IVT samples were then analysed for β-galactosidase activity, β-galactosidase amounts or used as an input for a sucrose cushion centrifugation (264,000 g, 2 hours, 4 °C) through a RNC low-salt sucrose cushion. For comparison to the activity of in vivo and IVT-produced β-galactosidase (Fig 1c), purified enzyme was titrated into an IVT mixture in the absence of plasmid DNA.

### Light scattering

Scattering was measured at 320 nm using a UV/visible spectrophotometer (V-760, Jasco) equipped with a temperature-controlled cuvette chamber. The unfolding reactions contained 10 µM β-gal (or 50 µM), 8 M urea, 100 mM sodium phosphate pH 7.5, 1 mM EDTA and 10 mM DTT. The unfolding reactions were diluted 10-fold into buffer containing 100 mM sodium phosphate pH 7.5, 1 mM MgCl_2_, 5 mM DTT, and either 0.7 or 0 M urea. The reactions were mixed and transferred to a cuvette held at 10 or 25 °C.

### Native PAGE

30 µl of refolded or native control reactions containing 1 µM β-gal were mixed with 10 µl of 4X native loading buffer containing 0.4 M Tris-HCl pH 8.0, 40 % (v/v) glycerol, 0.01 % (w/v) bromophenol blue, and snap frozen in liquid nitrogen. The samples were thawed and loaded immediately into cold 7% NuPage Tris-Acetate gels (ThermoFisher). The gel tank and the running buffer (24 mM Tris base, 191 mM glycine) were cooled for several hours in the cold room before loading the samples. For activity staining, the gels were incubated in a solution containing 50 mM Tris-HCl pH 7.0, 150 mM NaCl, 10 mM MgCl_2_ and 0.1 mg ml^-1^ X-gal, at 23 °C. For western blotting, the gel was incubated with 1x PBST for 5 min and the proteins were blotted to a PVDF membrane using the Trans-Blot Turbo system (Bio-Rad). The membrane was blocked using 5 % (w/v) skimmed milk power in PBST and incubated with the β-gal antibody (SC-65670, Santa Cruz Biotechnology) overnight at 4 °C. The membrane was washed and incubated with anti-mouse HRP secondary antibody (ab205719, Abcam) and developed using SuperSignal West Pico PLUS chemiluminescent substrate (ThermoFisher). Imaging was done using Amersham ImageQuant 800 (Cytiva). The gels were stained using Quick Coomassie Stain (Neo-Biotech).

### Limited proteolysis

RNCs were diluted to 0.5 μM in RNC low-salt buffer and cooled to 4 °C. Subsequently a reference aliquot was removed before proteolysis was initiated by addition of Proteinase K to a final concentration of 2.5 ng/μL. Aliquots of the proteolysis reactions were removed and quenched at different times by 1:1 mixing with 5 mM PMSF in RNC low-salt buffer. Quenched sample were then resolved by SDS-PAGE.

### Negative stain EM

Full-length β-gal or β-gal truncation 1-725 purified from Δβ-gal cells were diluted to 1 μM in RNC low-salt buffer. 4 µL were applied to copper grids (EM Resolutions, C300Cu100) coated with a continuous carbon only support films (15-20 nm) that had been glow discharged (25 mA, 60s) in a GloQube Plus Glow Discharge System (Quorum Technologies). After 1-2 minutes of incubation, excess sample was removed with filter paper (Whatman, 1001-090). Next, 40 µL of 2% uranyl acetate was applied three times for 20 s. Excess stain was removed with filter paper after each application and finally grids were air-dried. Micrographs were collected on a FEI Tecnai 12 Spirit 120 kV TEM with magnification 67,000x in a low-dose mode.

### Cryo-EM sample preparation

4 µl of purified RNC_1-510_, prepared as described above (50 mM HEPES pH 7.5, 12 mM MgOAc2, 100 mM KOAc, 1 mM DTT, RiboLock RNase inhibitor and 0.1% octyl glucoside), was deposited on 45 nm gold film 1.2/1.3 hole pattern 300 mesh grids (au-flat^TM^, EMS) glow-discharged in air for 60 s at 40 mA in an Emitech K100X glow discharge unit to render them hydrophilic, then blotted for 6 seconds using a Vitrobot Mark IV (FEI/Thermo Fisher Scientific) at 4 °C and 100% humidity and flash frozen in liquid ethane.

### Cryo-EM data collection and single particle analysis

In total, 9,384 micrographs were collected using Titan Krios cryo-EM (Thermo Fisher Scientific) operated at 300 kV, equipped with a Falcon4i detector. Images were collected using EPU software (Thermo Fisher Scientific) with a total of 1935 EER frames and nominal magnification of 75,000× (1.08 Å per pixel). The electron dose was 29.8 electrons per Å^2^ with dose per frame being 1.192 electrons per Å^2^. The defocus range was −1.3 to −2.8 μm. Beam-induced motion correction and per-frame radiation damage weighting with 5 by 5 patches were performed using MotionCor2 algorithms implemented in RELION^65^. CTF estimation was done on motion corrected micrographs with CTFFIND-4.1^66^. Micrographs with estimated resolution of 9 Å or higher were retained, resulting in 58 discarded micrographs. Initial particle picking was done using TOPAZ^67^, on a smaller micrograph subset (348 micrographs) with particle diameter of 280. 25,827 particles were selected and used to train a TOPAZ picking model. Using the full dataset, 4,974,533 particles were picked. Particle coordinates were extracted using a square box size of 320 pixels with 2-fold binning. Extracted particles were subjected to two rounds of 2D classification in RELION, with the first round ignoring CTFs until first peak. 835,178 particles were selected for a consensus 3D refinement, using an *ab-initio* map from CryoSPARC4.6^68^ as a reference. From the resulting consensus refinement at 4.37 Å resolution, a solvent mask was created and used for 3D classification without alignment. Here, there were 3 classes, one with 50S-only particles (240,640 particles, 28.8%), one with poor resolution 70S (109,325 particles, 13.1%) and the largest class containing well-aligned 70S particles (485,213 particles, 58.1%). The latter class was further classified based on tRNA site occupancy. The particles were imported into CryoSPARC, where utilising an 80 Å spherical mask and 3D variability analyses^69^, 3 clusters of particles were identified. Classes containing both the A- and P- site tRNA were pooled together to a total of 389,472 particles (80.2%). These particles were then refined in CryoSPARC via Non-Uniform Refinement^70^ and transferred back to RELION via pyem0.65. The particles were re-extracted without binning and subjected to 3D refinement in RELION followed by iterative rounds of CTF refinement where anisotropic magnification, 4th order aberrations and trefoil were estimated and followed by per-particle defocus and per-micrograph astigmatism fitting. Particles were then subjected to per-particle motion correction by Bayesian polishing. A final refinement resulted in a 2.86 Å-resolution reconstruction. The map was then sharpened using a B-factor of 61.45 during post-processing in RELION, as shown in Fig S9, otherwise the unsharpened map has been used. Associated figures were created using UCSF ChimeraX version1.9^71^.

### Model building and real-space refinement

Empty ribosome structures (PDB: 7K00^72^; 8QOA^37^) were used as initial models for atomic model building. These models were rigid-body fitted into map using UCSF ChimeraX. Then, each chain was real-space-refined in ISOLDE^73^. The nascent chain was built *de novo* based on RNC_1-510_ sequence, first fitting residues of the amphipathic helix _466_ANHDALYRWIKSV_478_ and stalling sequence WWWPRIRGP as these had the best-defined side-chains. The atomic models were subjected to multiple rounds of real-space refinement in Phenix version 1.21.2^74^ and ISOLDE was used to manually fix all outliers and bad geometries. Validation of the models was carried out using Phenix, wwwPDB Validation, and the Molprobity server^75^ (http://molprobity.biochem.duke.edu/).

### Circular dichroism spectroscopy

Peptide Ac-ANHDALYRWIKSV-NH_2_ synthesized in-house was resuspended in water and diluted to 0.15 mg/mL in 20 mM sodium phosphate buffer, pH 7 prior to circular dichroism measurement at 20 °C. The average of 25 scans (range = 190 to 260 nm, bandwidth = 2 nm, increment = 0.2 nm) was recorded and corrected using a buffer blank. The data were subsequently deconvoluted to estimate the relative secondary structure content using ContinLL, SELCON3, and CDSSTR methods^76^. The average value obtained from the three deconvolution methods was reported.

### Quantification and statistical analysis

Band intensities on immunoblots and SDS-PAGE gels were quantified in Fiji^64^. All statistical analyses and curve fitting were performed in GraphPad Prism 10. Hydrophobicity values of solved protein structures were calculated using ChimeraX Molecular Lipophilicity Potential. Hydrophobic clusters composed of Ile, Leu and Val were identified using the ProteinTools server^77^ (https://proteintools.uni-bayreuth.de/). The residues constituting the long and activating interface were identified using the PDBePISA server^78^.

### Pulse-labelled HDX-MS analysis of β-gal refolding

The refolding reaction was set-up as described above. At different times, reactions were exchanged into the deuteration buffer using spin desalting columns (Micro Bio-Spin 6, Bio-Rad). The columns were equilibrated four times with 500 µl of deuteration buffer composed of 10 mM sodium phosphate (pH_read_ = 7.01 at 23 °C), 1 mM MgCl_2_, 5 mM DTT, 1.5 M urea (99.9% D2O). The deuteration buffer was prepared by mixing the stock solutions of the components which were prepared in D_2_O and the final pH_read_ was 7.48 at 23 °C. Undeuterated control reaction was exchanged into a solution prepared with water but identical in composition to deuteration buffer. 50 µl of the refolding reaction was pipetted into the spin columns and spun at 1000 ×g for 1 min at 23 °C. The exchange was immediately quenched by mixing 45 µl of the deuteration reaction with 15 µl of quench buffer (1 M orthophosphoric acid with pH adjusted to 2.12 using NaOH), and the tube was flash frozen in liquid nitrogen and stored at -70 °C. Triplicate independent refolding reactions were performed and these results are reported in the manuscript. The experimental outcome was validated by performing a biological replicate experiment using an independent protein purification where fewer time points of refolding were sampled.

The samples were thawed and 50 µl was immediately injected into an Acquity UPLC M-class system (Waters) equipped with an HDX manager. The injection order was randomized. The protein was pumped at 200 μL min^-1^ for 2 min over an Enzymate BEH pepsin column (5 μm, 2.1 mm x 30 mm, Waters) held at 20 °C. The peptides were trapped and desalted on an Acquity UPLC BEH C18 Van-guard trap column (1.7 μm, 2.1 mm x 5 mm, Waters) held at 0 °C. The peptides were eluted at 40 µl min^-1^ and separated on an Acquity UPLC BEH C18 column (1.7 μm, 100 mm x 1 mm, Waters) held at 0 °C, using a 3-35% gradient of acetonitrile in 0.1 % (v/v) formic acid over 16 min. The peptides were detected using a SYNAPT G2-Si HDMS^E^ instrument in ion mobility mode, acquiring in positive ion mode over a range of 50 to 2,000 m/z. The electrospray ionization source temperature was 80°C and the spray voltage was 3.0 kV. The lock mass calibration was performed using [Glu^1^]-Fibrinopeptide B (50 fmol/μL, Merck, F3261).

To prepare the maximally deuterated control, 1 µM β-gal in 0.1 % formic acid was digested for 5 min at 25 °C using pepsin beads. The beads were separated using a spin filter, and 45 µl of the digest was dried in a speed vac. The peptides were resuspended in 45 µl of D_2_O and incubated for 18 h at 35 °C. This was then mixed with 15 µl of the quench buffer and injected immediately into the mass spectrometer as described above.

The peptides in undeuterated β-gal were searched using Protein Lynx Global Server (version 3.0.3, Waters) search engine. In the workflow parameters, a database consisting of β-gal sequence, non-specific cleavage and oxidation of methionine as variable modification were specified. A low energy threshold of 135 counts and elevated energy threshold of 30 counts were specified in the processing parameters. The peptides identified by Protein Lynx Global Server were input into DynamX (version 3.0.0, Waters) and filtered using the settings minimum products per amino acid of 0.2 and minimum consecutive products of 2. The spectra were manually inspected to rectify incorrect peak assignments and peptides with poor quality spectra were rejected.

DynamX analysis was exported as a cluster file. The mass of the peptide was calculated using the formula PepMass = Z × (Center – Proton Mass). Relative deuterium uptake of a peptide was calculated by subtracting the average mass of triplicate values for the undeuterated peptide from the mass of the deuterated peptide. For each peptide, the relative uptake of refolded sample (29 h of refolding) was subtracted from unfolded sample (0.06 h, first time point sampled for the refolded protein). Peptides experiencing an overall change in mass less than 0.5 Da were given a half-life of 0 min. These were typically located in exposed loops which do not change in deuterium uptake during refolding. For the remaining peptides, data describing relative uptake versus time were exported into Graphpad Prism, and fit to one-phase decay equation to generate folding half-times. Per-residue half-time was calculated by averaging the values from overlapping peptides. All plotted uptake data and a summary of experimental conditions^79^ are shown in Data S1. The MS data have been deposited to the ProteomeXchange Consortium via the PRIDE partner repository with the dataset identifier PXD063232^80^.

### Equilibrium HDX-MS analysis of RNCs and β-gal truncations

HDX-MS was conducted as described previously^58^ with some modifications. In short, stock samples were prepared in RNC low-salt buffer at final 5-6 μM for RNCs and empty ribosomes (NEB, P0763S), and at final 15-20 μM for C-terminally truncated or full-length β-galactosidase. Deuterium labelling was initiated by mixing 3 μL of stock samples with 27 μL deuteration buffer (10 mM HEPES-NaOH, pH 7.1, 30 mM KOAc, 12 mM Mg(OAc)_2_, 1 mM DTT, RiboLock RNase inhibitor (ThermoScientific), 97% D_2_O) at 25 °C. After labelling at 25 °C for 10 or 100 seconds, the reaction was quenched with an equal volume (30 μL) of ice-cold quench buffer (100 mM sodium phosphate, pH 1.4, 4 M guanidium hydrochloride, 10 mM TCEP). The pH after quenching was 2.5. Digestion was initiated by addition of 20 μL pepsin-agarose 50% v/v bead slurry (prepared in-house as described previously^58^) equilibrated in 0.1% formic acid. Following 100 s of digestion at 10 °C with rapid mixing every 30 seconds, the sample was centrifuged (13,000 g, 15 s, 0 °C) through 0.22 μm PVDF filters (Millipore, UFC30GVNB). The flow through was immediately snap-frozen in liquid nitrogen for short-term storage. The same protocol was followed to prepare undeuterated controls, except the deuteration buffer was replaced by a H-based buffer (10 mM HEPES-NaOH in H_2_O, pH 7.5, 30 mM KOAc, 12 mM Mg(OAc)_2_, 1 mM DTT, RiboLock RNase inhibitor).

Frozen samples were rapidly thawed and immediately injected into an Acquity UPLC M-class system with the cooling chamber containing the chromatographic columns kept at 0 ± 0.2 °C throughout data collection. Peptic peptides were trapped and desalted for 4 minutes (200 μL/min) on a 2.1 mm X 5 mm, C4 trap column (Acquity BEH C4 Van-guard pre-column, 1.7 μm, Waters, 186004623) then separated on a reverse phase Acquity UPLC HSS T3 column (1.8 μm, 1 mm X 50 mm, Waters, 186003535) at a flow rate of 90 μL/min. Peptides were eluted over 25 minutes using a 3-30% gradient of acetonitrile in 0.1% formic acid. Analysis was performed using a Waters Synapt G2Si HDMS^E^ instrument in ion mobility mode, acquiring in positive ion mode over a range of 50 to 2,000 m/z with the conventional electrospray ionisation source. Calibration of the mass spectrometer was achieved using [Glu^1^]-Fibrinopeptide B (50 fmol/μL, Merck, F3261) and the instrument was operated at a source temperature of 80 °C with the capillary set to 3 kV.

MS^E^ data were processed using Protein Lynx Global Server (PLGS, Waters) to identify peptides in the undeuterated control samples using information from a non-specific cleavage of a database containing sequences of *E. coli* β-galactosidase, Trigger Factor, 70S ribosomal proteins as well as porcine pepsin. PLGS search was performed using energy thresholds of low = 135 counts and elevated = 30 counts. Peptides identified by PLGS were subsequently filtered and processed in DynamX (Waters) with filters of minimum products per amino acid of 0.05 and minimum consecutive products of 1. All spectra were manually inspected, and poor-quality assignments were removed. Any peptides assigned to nascent chains that were also detected in empty ribosome control samples were removed. Additionally, peptides assigned to β-galactosidase in nascent chains but not in isolated β-galactosidase, were also removed. Relative deuterium uptake in Da was calculated by subtracting the centroid mass of undeuterated peptides from those of deuterated peptides. Determined mean values of deuterium uptake are relative as they are not corrected for back-exchange. All HDX-MS experiments were performed in technical triplicates for each sample at each deuteration time and used to calculate the reported mean. In some cases, we also performed biological replicates using independent purifications. All plotted uptake data, peptide coverage maps, and a summary of experimental conditions^79^ are shown in Data S2. The MS data have been deposited to the ProteomeXchange Consortium via the PRIDE partner repository with the dataset identifier PXD060082^80^.

### Selective ribosome profiling (SeRP)

lacZ-mEYFP under the control of the IPTG inducible tac promoter was integrated in the E. coli MC4100 genome at the λ-phage attachment site^81^. Exponentially growing cells were harvested at OD600 = 0.45 – 0.5 by rapid-filtration and flash-freezing in liquid nitrogen. Expression of lacZ-mEYFP was induced by 10 μM IPTG (f.c.) for 15 min before cell harvest. Cell pellets were lysed under cryogenic conditions by mixer milling (2 min, 30 Hz) with SeRP lysis buffer (50 mM HEPES KOH pH 7.0, 100 mM NaCl, 10 mM MgCl2, 5 mM CaCl2, 50 μg/ml chloramphenicol, 1 mM PMSF, cOmplete EDTA-free protease inhibitor (Roche)). Upon thawing on ice, RP lysis buffer containing DNaseI was added to the lysate and the lysate was cleared by centrifugation (20 000 xg, 10 min, 4 °C). The supernatant was digested with self-made micrococcal nuclease (MNaseI) for 1 h at 25 °C, quenched by EGTA (f.c. 6 mM). For the total translatome, ribosomes from one-fifth of the lysate were purified by sucrose density centrifugation (20 mM Tris-HCl pH 8.0, 100 mM NH4Cl, 10 mM MgCl2, 50 μg/ml chloramphenicol) and ribosome-protected mRNA-footprints were isolated by acid phenol extraction as previously described^82^. For the selected translatome, the remaining lysate was incubated with GFP-binder coupled sepharose beads (800 μl 1:1 suspension) for 1 h at 4 °C by end-to-end rotation. Beads were washed three times with wash buffer (50 mM HEPES KOH pH 7.0, 100 mM NaCl, 10 mM MgCl2, 50 μg/ml chloramphenicol, 1 mM PMSF, cOmplete EDTA-free protease inhibitor (Roche), 0.02% (v/v) Tween-20) for 5 min at 4 °C. Ribosome protected footprints were extracted by acid phenol extraction. cDNA libraries for deep sequencing analysis were generated as described previously^82^ and sequenced on a HiSeq 2000 v4 (Illumina).

### Disome Selective Profiling (DiSP)

E. coli MG1655 cells were grown at 37 °C in EZ RDM with lactose as the sole carbon source and harvested by rapid filtration and flash-freezing in liquid nitrogen at OD600 = 0.45 – 0.5. Cells were lysed in the presence of DiSP lysis buffer (50 mM HEPES KOH pH 7.0, 100 mM NaCl, 10 mM MgCl_2_, 5 mM CaCl_2_, 50 μg/ml chloramphenicol, 1 mM PMSF, cOmplete EDTA-free protease inhibitor (Roche), 0.4% (v/v) Triton X-100, 0.1% Nonidet P40 Substitute) under cryogenic conditions by mixer milling (2 min, 30 Hz). Upon thawing 750 uL DiSP lysis buffer containing DNase I was added and the lysate was cleared by centrifugation (20 000 xg, 10 min, 4 °C). The supernatant was digested with self-made micrococcal nuclease for 10 min at 25 °C, quenched by EGTA (f.c. 6 mM). 250 μg total RNA was layered over a 5-45% sucrose gradient (50 mM HEPES-KOH pH 7.0, 500 mM KCl, 10 mM MgCl_2_, 50 μg/mL chloramphenicol) and separated by ultracentrifugation for 3.5 h at 35.000 rpm, 4 °C. Monosome and disome fractions were isolated and the RNA extracted by acid phenol. cDNA libraries for deep sequencing analysis were generated as described previously^83^ and sequenced on a NextSeq 550 (Illumina).

### Ribosome profiling data analysis

HiSeq 2000 SeRP data was processed and analyzed as described previously^82^. NextSeq550 DiSP data was processed and analyzed as described previously^20^.

## DATA AVAILABILITY

The sequencing data discussed in this publication have been deposited in NCBI’s Gene Expression Omnibus and are accessible through GEO Series accession number GSE291748. Data analysis of ribosome profiling datasets was performed with RiboSeqTools (available at https://github.com/ilia-kats/RiboSeqTools).

All mass spectrometry data have been deposited to the ProteomeXchange Consortium via the PRIDE partner repository^84^ with the following dataset identifiers:

HDX-MS analysis of β-gal refolding: PXD063232

HDX-MS analysis of β-gal truncations and RNCs: PXD060082

The cryoEM map has been deposited in the Electron Microscopy Data Bank (EMDB), and the coordinates have been deposited in the PDB under the following accession numbers: EMD-53553 and PDB: 9R3A.

## SUPPLEMENTARY FIGURES AND LEGENDS

**Fig. S1.**
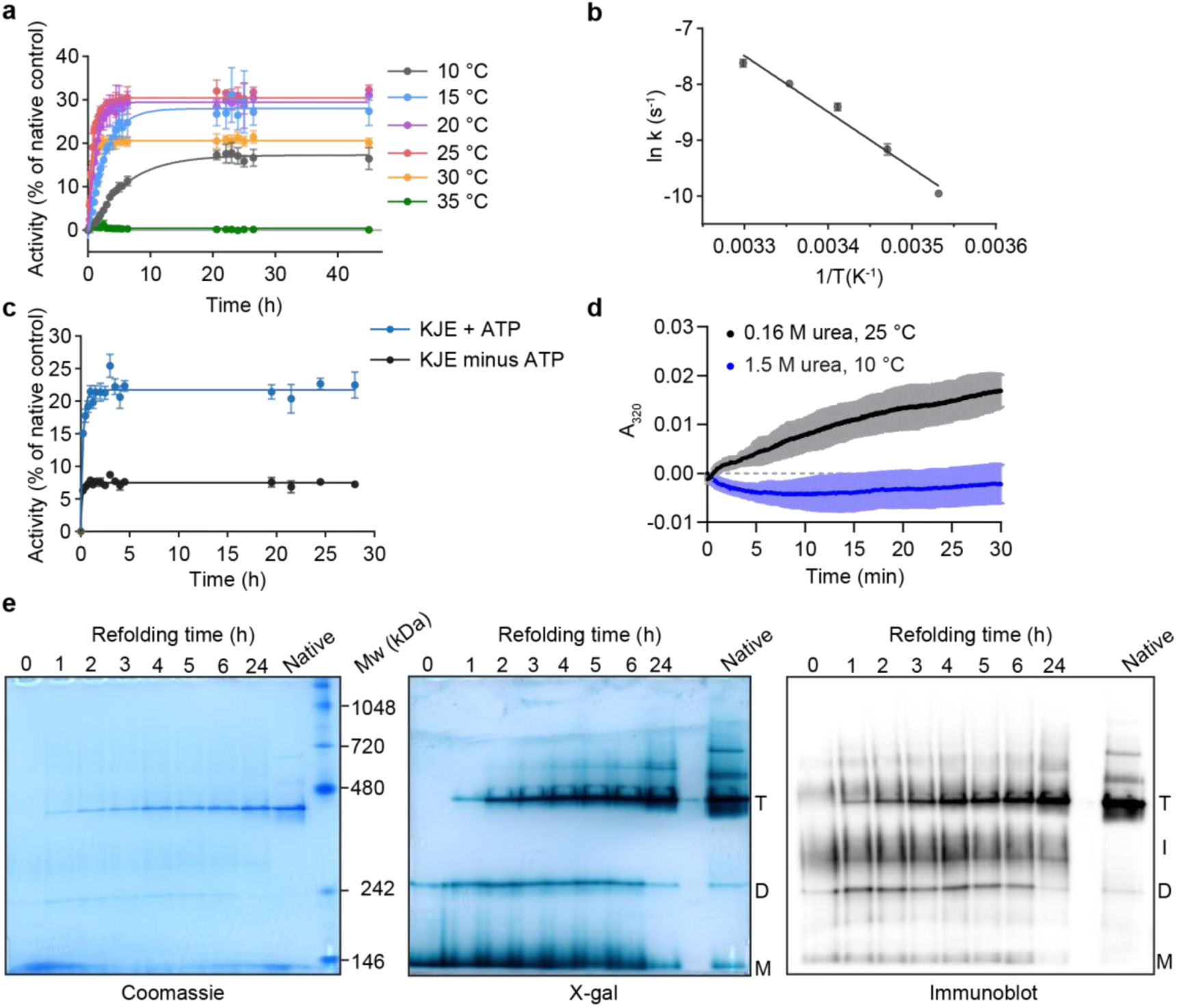
β-gal refolding. a, β-gal refolding upon dilution from 8 M urea into buffer at different temperatures, assayed by recovery of enzyme activity. Data were fit to a single exponential function. Error bars represent s.d. (n = 3). **b,** Arrhenius plot showing temperature dependence of β-gal refolding kinetics as measured in (**a**). *E*_a_ = 84±1 kJ.mol^-1^. Error bars represent s.d. (n = 3). **c,** Influence of Hsp70 chaperone system on β-gal refolding. Refolding was measured as in (a), at 37 °C. The reaction was supplemented with purified DnaK, DnaJ and GrpE, with or without ATP. Data were fit to a single exponential function, giving t_½_ = 11 min (95% CI, 8 – 14 min) in the presence of ATP. Error bars represent s.d. (n = 3). **d,** Aggregation during β-gal refolding. Turbidity was monitored by absorbance at 320 nm. Refolding under optimal conditions as in Fig. 1b (10 °C, final urea concentration of 1.5 M) did not result in detectable turbidity. Turbidity could be induced by raising the temperature of the reaction to 25 °C and reducing the final urea concentration to 0.16 M). The shaded region represents s.d. (n = 3). **e**, Oligomeric state of β-gal during refolding, analysed using native-PAGE. Replicate gels were stained with Coomassie, immunoblotted for β-gal, or stained with the β-gal substrate X-gal. Bands corresponding to the tetramer (T), dimer (D), monomer (M) or intermediate (I) are indicated. Natively folded β-gal purified from *E.coli* is included as a control.

**Fig. S2.**
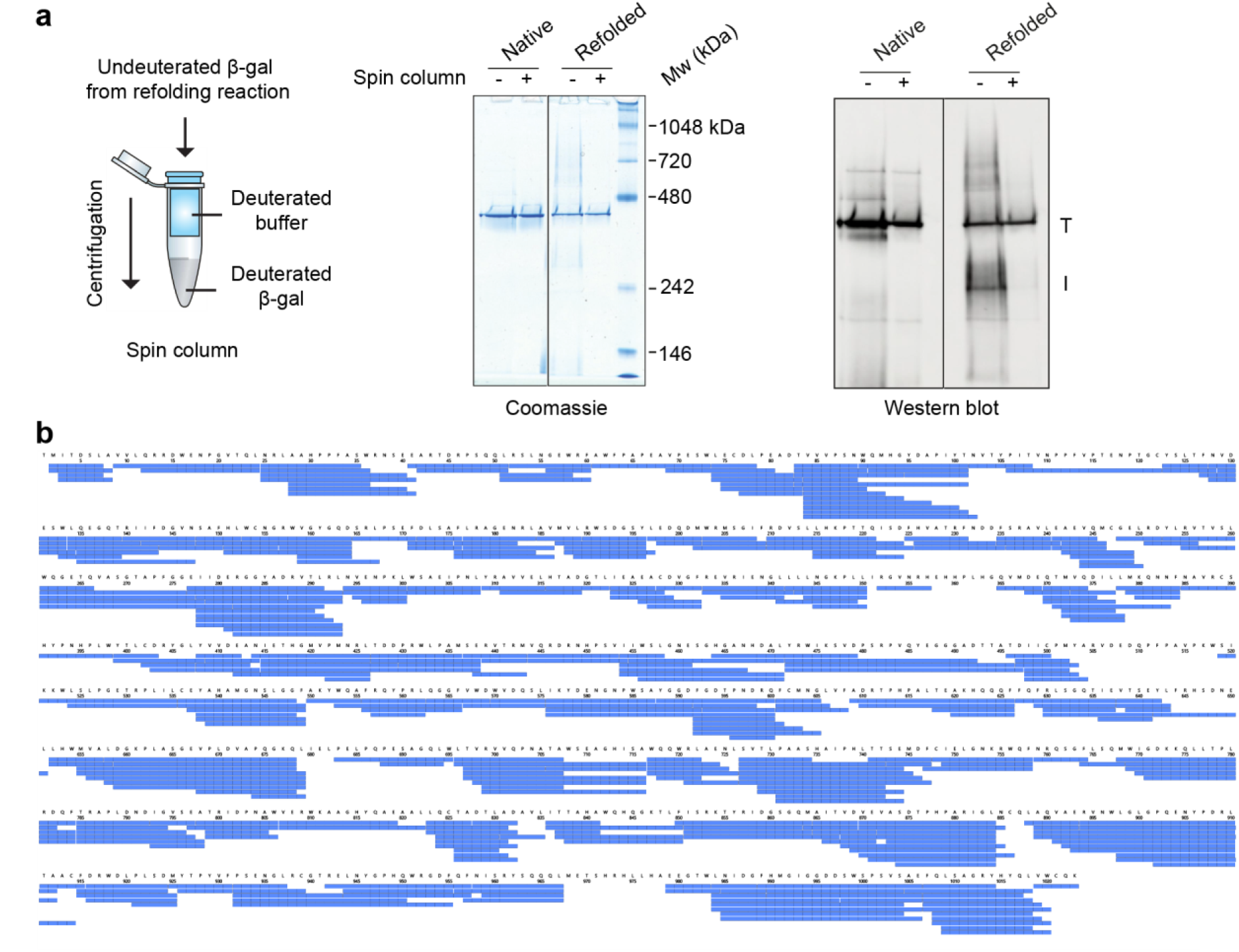
HDX-MS analysis of β-gal refolding. a, Spin-column buffer exchange depletes non-native oligomers from the refolding reaction. Native, or refolded β-gal was resolved by native-PAGE, before (-) or after (+) centrifugation through a spin column. Duplicate gels were stained with Coomassie or immunoblotted for β-gal. Bands corresponding to the tetramer (T) and oligomeric intermediate (I) are indicated. **b**, HDX-MS peptide coverage of β-gal. 400 peptides (blue bars) were followed during the refolding reaction, covering 96.3% of the sequence.

**Fig. S3.**
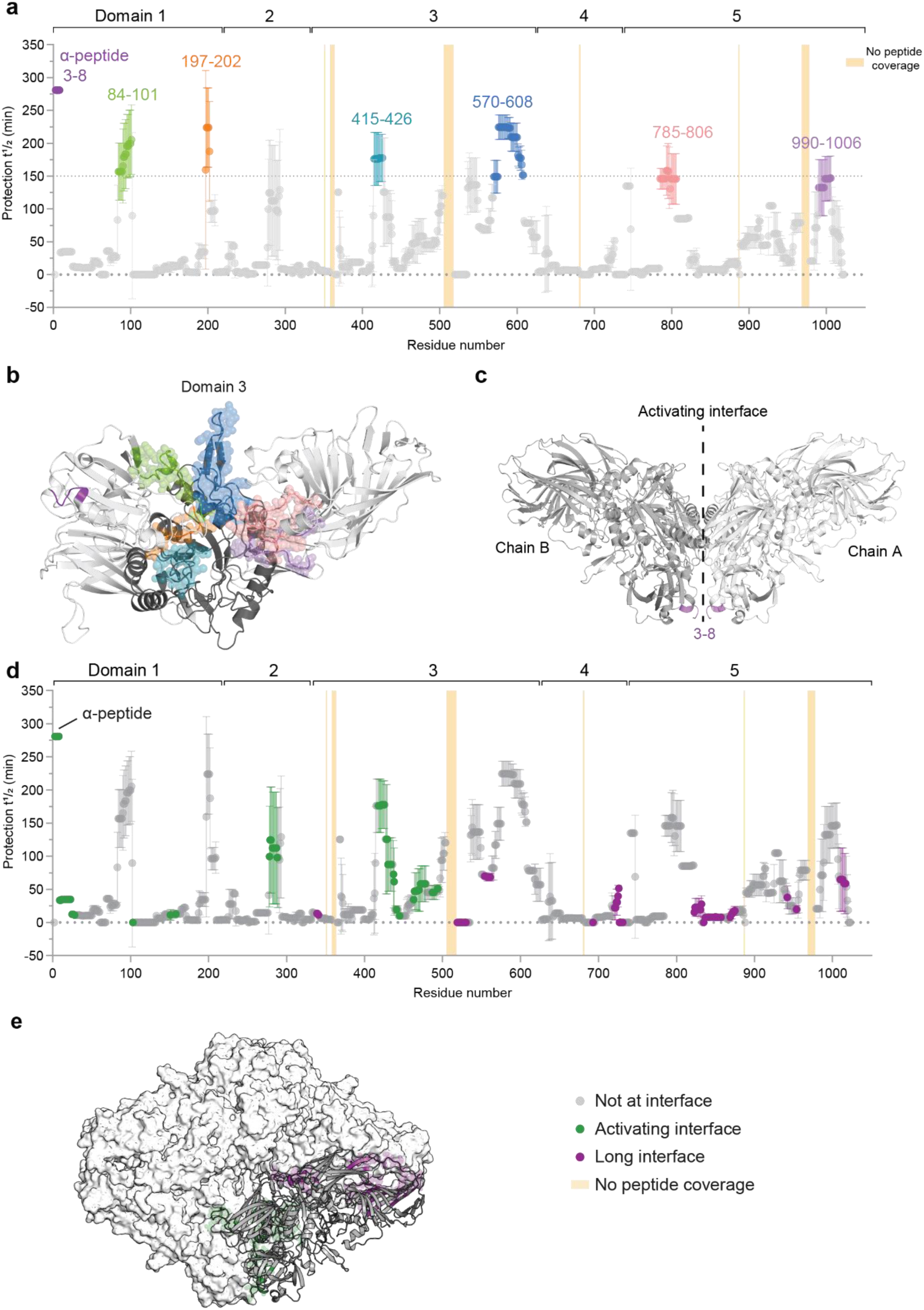
Slow steps during β-gal refolding. a, Residue-resolved refolding half-times are plotted as in Fig. 2d, and residues with t_½_ ≥ 150 min are indicated. **b**, Slow-folding regions cluster in native β-gal. Residues highlighted in (**a**) are coloured on the structure of the β-gal monomer. **c**, Peptide 3-8, forming part of the α-peptide^27^, is at the activating interface. Residues 3-8 are coloured purple on the native structure of β-gal with two subunits shown. **d,** Residue-resolved refolding half-times are plotted as in Fig. 2d, with residues at the long and activating interfaces coloured purple and green, respectively. Error bars represent s.d. (n = 3). **e,** Structure of β-gal tetramer, coloured as in (**d**).

**Fig. S4.**
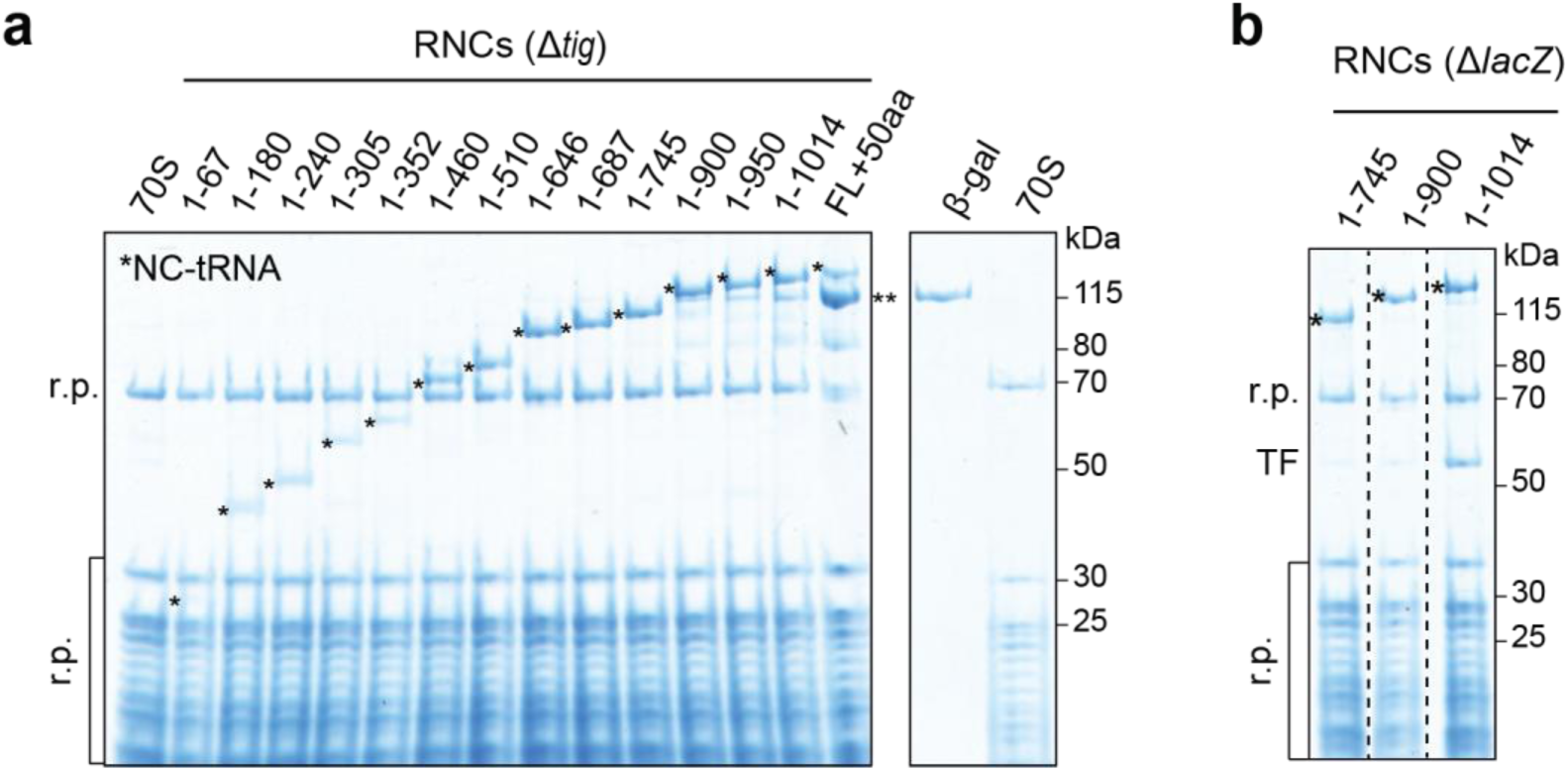
β-gal ribosome-nascent chain complexes (RNCs). a,b, Coomassie-stained gel of RNCs purified from *E.coli* lacking endogenous Trigger factor (Δ*tig*) or β-gal (Δ*lacZ*). Ribosomal proteins (r.p.), Trigger factor (TF), the nascent chain linked to its peptidyl-tRNA (*) and co-purifying full-length β-gal (**) are indicated. Purified β-gal and empty 70S ribosomes are shown alongside for reference.

**Fig. S5.**
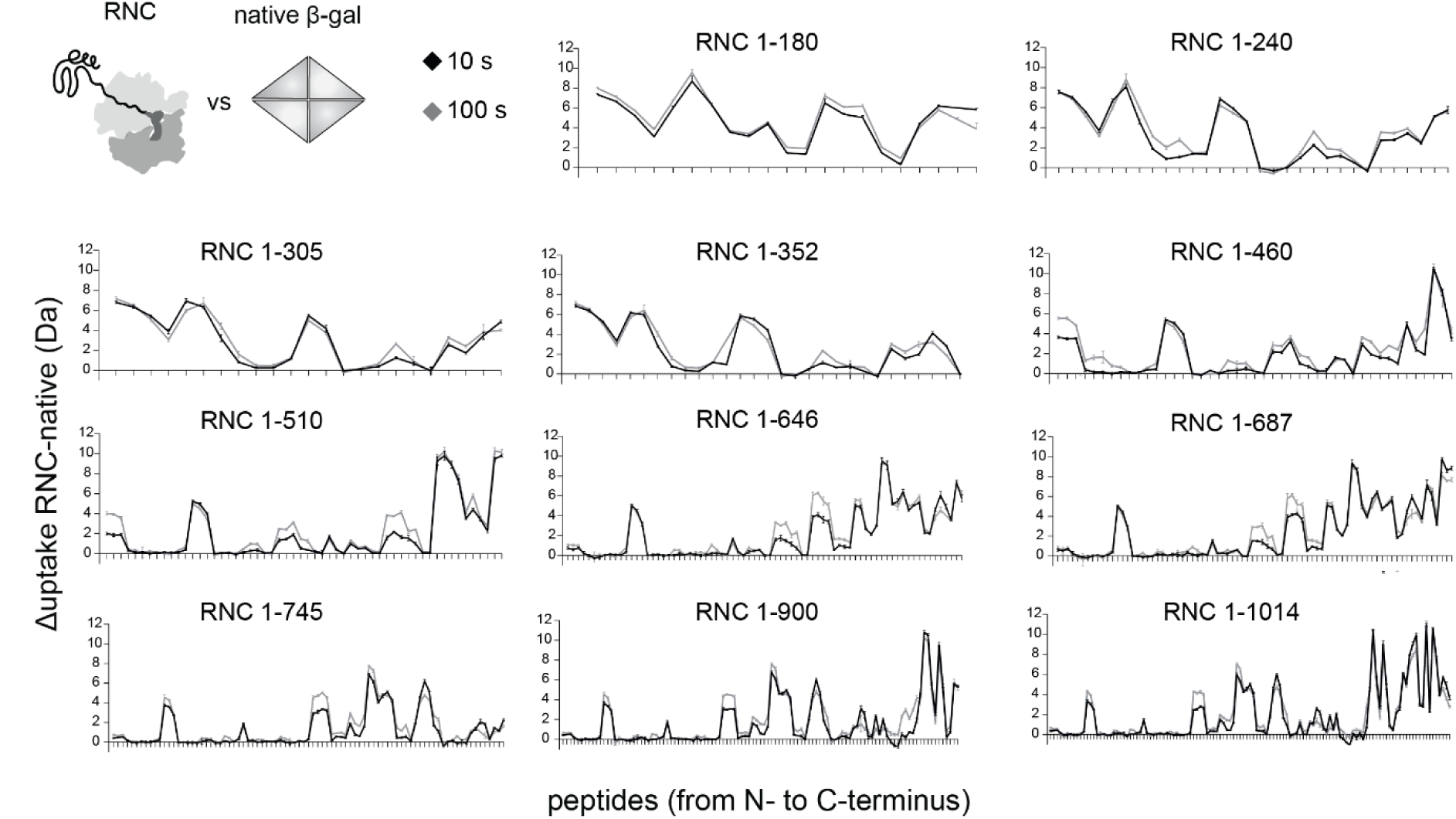
HDX-MS analysis of RNCs. Deuterium uptake of β-gal NCs relative to native β-gal. Difference in deuterium uptake, after 10 s (black) or 100 s (grey) deuteration, between NC peptides and full-length native β-gal. All RNCs between 1-180 and 1-687 were purified from Δ*tig* cells and RNCs 1-745, 1-900, and 1-1014 were purified from Δ*lacZ* cells. Higher values indicate more deuteration of RNCs relative to native β-gal. Error bars represent s.d. (n = 3).

**Fig S6.**
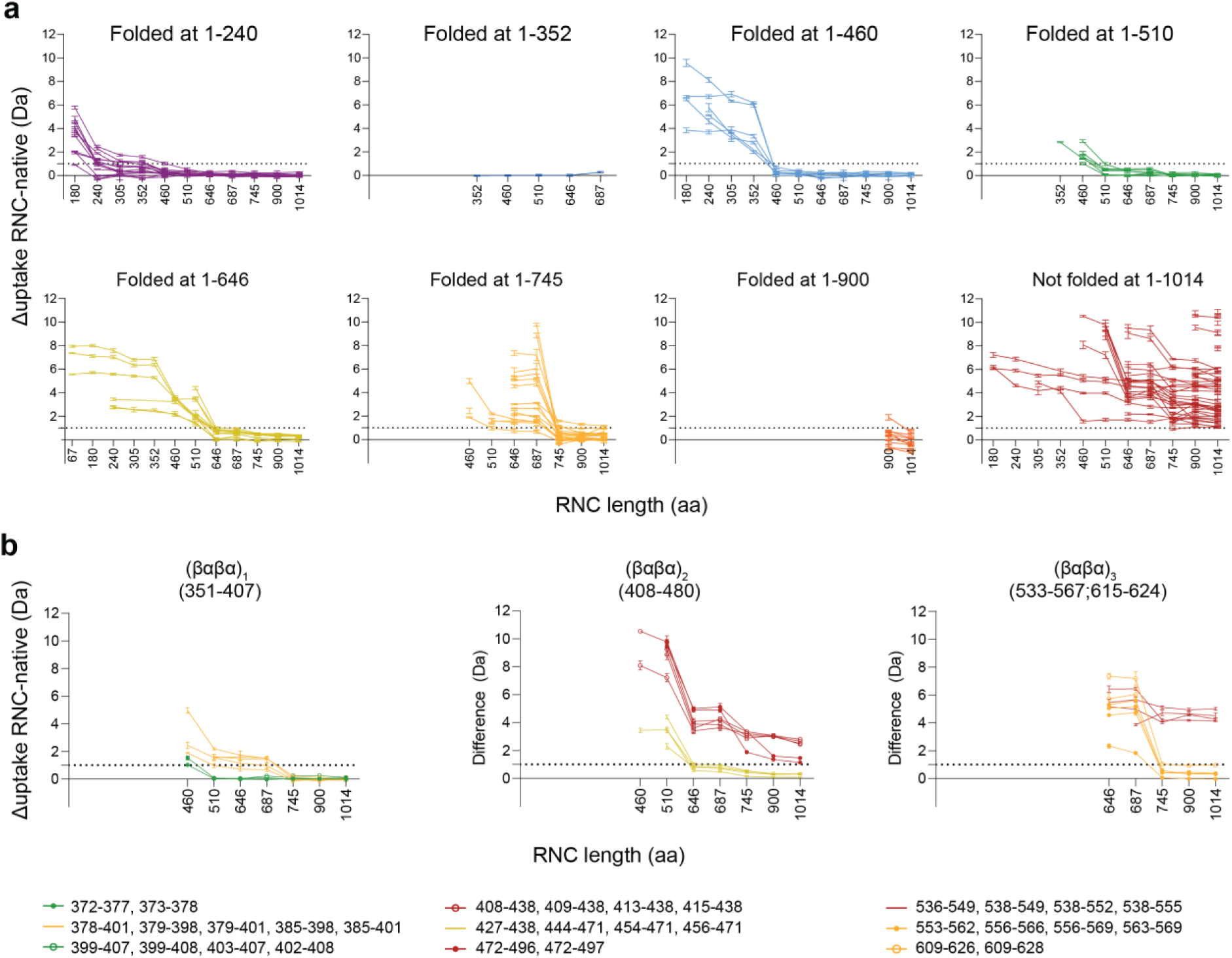
β-gal cotranslational foldons. a, Deuterium uptake of β-gal peptides as a function of NC length. Plots show the difference in deuterium uptake, after 10 s deuteration, between peptides in indicated RNCs and full-length native β-gal. Peptides are clustered according to the NC length at which their level of deuterium uptake plateaus (i.e. when uptake changes <1 Da between successive RNCs). Overlapping peptides are clustered together. Error bars represent s.d. (n = 3). **b**, as in (**a**), for different βαβα in domain 3.

**Fig S7.**
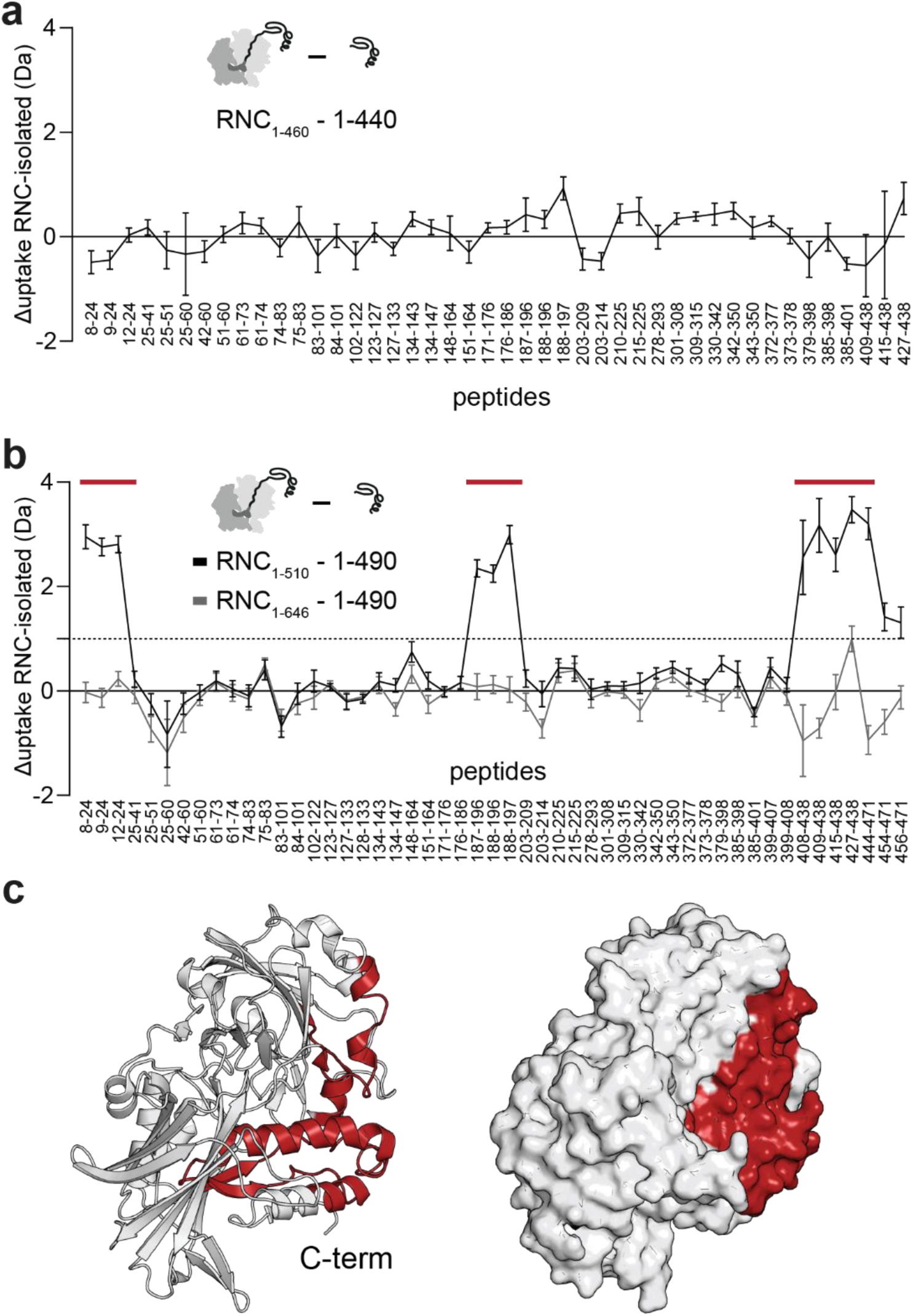
Destabilisation of β-gal on the ribosome during domain 3 synthesis. a, Difference in deuterium uptake, after 100 s deuteration, between truncated β-gal (1-440) and the corresponding RNC_1-460_. Error bars represent s.d. (n = 3). **b,** As in (**a**), showing the difference between truncated β-gal (1-490) and RNC_1-510_ or RNC_1-646_. Error bars represent s.d. (n = 3). **c**, Regions that are deprotected in RNC_1-510_ relative to 1-490 are indicated in red, and coloured on the native structure of residues 1-490 (PDB: 6CVM^85^).

**Fig. S8.**
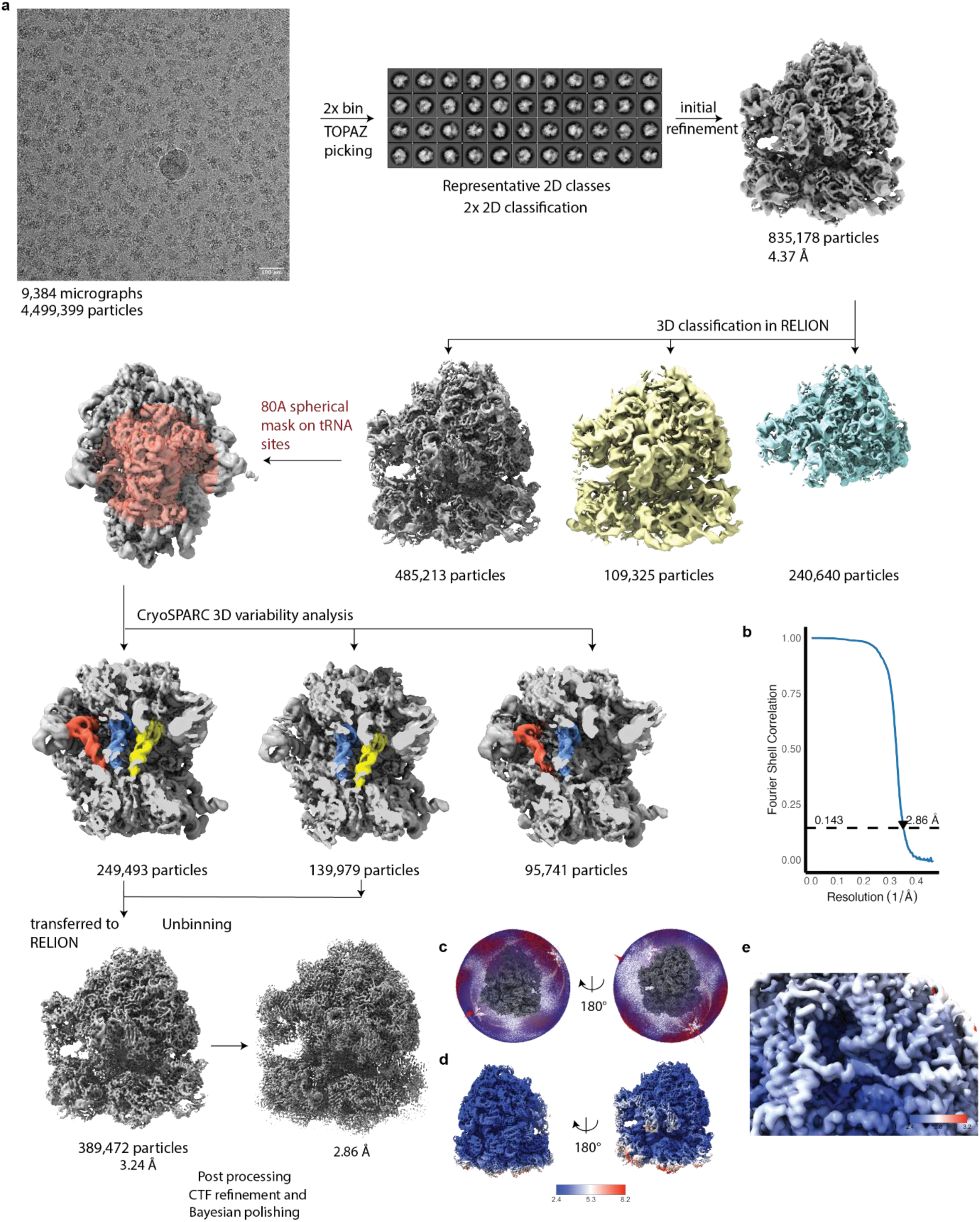
Cryo-EM data collection and single particle analysis of RNC_1-510_. a, Schematic diagram of the cryo-EM and single particle analysis workflow. On the left, a representative motion-corrected micrograph with scale bar representing 100 Å. 9,384 micrographs were collected at 1.08 Å/px and 4,499,399 particles were initially picked with TOPAZ and extracted with a box size of 320 Å. In the middle, representative high-quality 2D class averages, after iterative reference-free 2D classification and selection of the particles comprising the highest quality classes. On the right, a consensus 3D refinement of the thus selected 835,178 particles. These were then further classified without alignment. A single class comprised of high-resolution, well aligning 70S particles (grey) was selected and its underlying 485,213 particles (58,1% of the total) were imported into CryoSPARC and subjected to 3D variability analysis using an 80 Å spherical mask around tRNA sites (second row, left). Here, clusters containing A- and P- site tRNAs were pooled (80.2%) and subjected to non-uniform refinement in CryoSPARC (third row, first and second from the left). The output particles were transferred back to RELION 5, extracted without binning, and refined again (bottom row, left). These particles then underwent rounds of CTF refinement, Bayesian polishing and 3D refinement. The final structure is displayed after post-processing, at 2.86 Å overall resolution (bottom row, right). **b**, Fourier Shell Correlation (FSC) estimate by 0.143 criterion for RNC_1-510_^86^. **c**, Euler angle distribution plots for RNC_1-510_. **d**, Local resolution estimates for RNC_1-510_. **e**, Local resolution estimates for the area around the NC at the ribosome surface.

**Fig S9.**
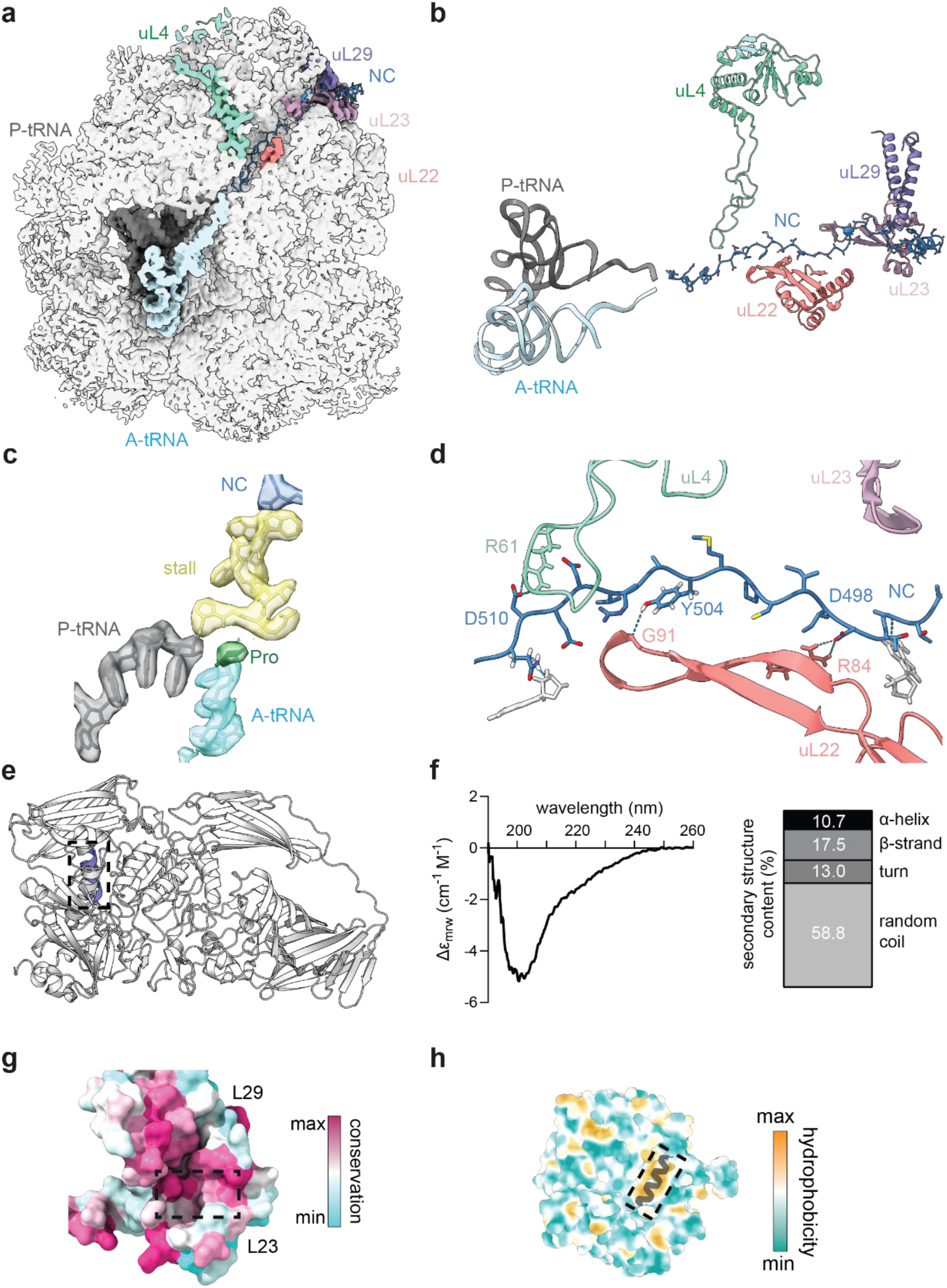
Structure and path of nascent β-gal on the ribosome. a, Path of the β-gal NC through the exit tunnel. Cross-section of the cryo-EM map of RNC_1-510_ with ribosomal proteins lining the exit tunnel highlighted, and the atomic model of the NC shown in cartoon representation. **b**, Molecular model of the NC and ribosomal proteins lining the exit tunnel. The NC follows the contour of L22 in the middle part of the tunnel, before pivoting towards L29/L23 close to the vestibule. **c**, Cryo-EM map (semi-transparent density) and molecular model (cartoon representation) for the stalling sequence (yellow), P-site tRNA (dark grey) and A-site tRNA (cyan). The proline amino acid on the A-tRNA is highlighted in green. **d**, Structure of the NC in the middle part of the exit tunnel, showing interactions with ribosomal proteins uL4 and uL22. **e,** Residues 466-478 fold into an α-helix in native β-gal. Structure of the β-gal monomer (PDB: 6CVM^85^) with residues _466_ANHDALYRWIKSV_478_ coloured blue and indicated with a dashed box. **f**, Secondary structure of isolated β-gal peptide. Left, circular dichroism spectrum for peptide Ac-ANHDALYRWIKSV-NH_2_ in 20 mM sodium phosphate buffer, pH 7. Right: estimated secondary structural content from spectral deconvolution. **g**, Evolutionary conservation of NC-interacting surface on L23 and L29. Conservation was calculated using ConSurf^87^. Only residues 1-88 of L23 are shown. The position occupied by the NC helix is indicated with a dashed box. **h**, β-gal helix occupies a hydrophobic groove in domain 3. Native structure of residues 1-490 of β-gal is shown in surface representation and coloured by hydrophobicity, except for residues 462-481 which are shown in cartoon representation and coloured black.

**Fig S10.**
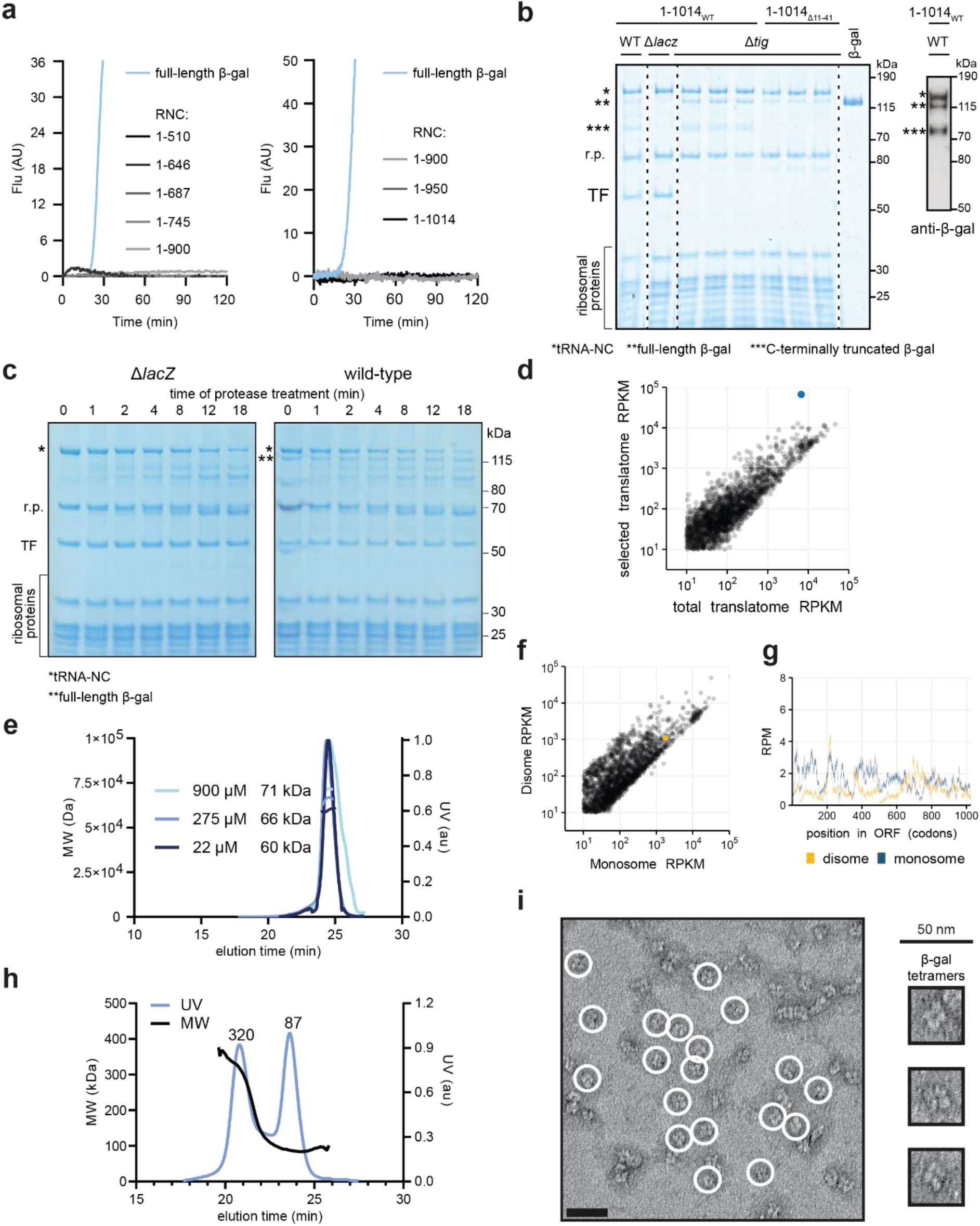
β-gal assembly. a, Isolated RNCs are inactive. Stalling constructs were expressed in PURE IVT reactions as in Fig. 1c. Enzyme activity was monitored in real-time by fluorescence upon cleavage of the substrate resorufin-β-D-glucopyranoside. Full-length β-gal (with a stop codon) was translated as a control. **b**, Copurification of endogenous full-length β-gal with RNC_1-1014_. Wild-type RNC_1-1014_ (RNC_1-1014WT_) or a variant with a disrupted activation interface^88^ (RNC_1-1014Δ11-41_) were purified from wild-type (WT), Δ*lacZ* or Δ*tig E. coli* and resolved by SDS-PAGE followed by Coomassie staining. The last lane contains purified full-length β-gal for reference. Bands corresponding to the NCs (*), copurifying full-length (**) or C-terminally truncated (***) β-gal, Trigger factor (TF) and ribosomal proteins (r.p.) are indicated. Right: immunoblot of RNC_1-1014WT_ purified from wild-type cells, probed using an antibody against the β-gal N-terminus (ab106567). **c**, Copurifying full-length β-gal is protease-resistant compared to the NC. RNC_1-1014_ purified from WT or Δ*lacZ* cells was treated with Proteinase K and resolved by SDS-PAGE followed by Coomassie staining. Bands corresponding to the NC (*) and to the full-length β-galactosidase (**) are indicated. **d,** β-gal pulldown specifically enriches *lacZ* mRNA. Comparison of the gene-specific footprint abundance in the β-galactosidase co-IP sample and the total translatome in RPKM (reads per kilobase per million mapped reads). The lacZ transcript is indicated in blue. **e**, 1-519 β-gal truncation is monomeric. Molecular weight was measured by SEC-MALS at different concentrations as indicated. Molecular weight (MW, left y-axis) and ultraviolet absorbance (UV, right y-axis) are plotted as a function of elution time. The theoretical mass of the 1-519 fragment is 59 kDa. **f**, *lacZ*-translating ribosomes are not enriched in the disome fraction. Comparison of the gene-specific footprint abundance in disome and monosome samples in RPKM (reads per kilobase per million mapped reads). The *lacZ* transcript is indicated in yellow. **g,** β-gal does not assemble via a co-co mechanism. Monosome (blue) and disome (yellow) footprint density along the lacZ open reading frame. RPM: reads per million. **h**, 1-725 β-gal truncation is partially tetrameric. SEC-MALS was performed as in (**d**), at a concentration of 30 μM. Two species of different molecular weights were detected, corresponding to monomeric (87 kDa) and tetrameric (320 kDa) forms. The theoretical mass of the 1-725 fragment is 83 kDa. **i**, Full-length β-gal forms tetramers rather than higher order assemblies as detected in Fig. 5e for the 1-725 fragment. Negative stain electron microscopy micrograph of purified full-length β-gal. The scale bar corresponds to 50 nm.

**Table S1.** Cryo-EM data collection, refinement and validation statistics.

|  |  |
| --- | --- |
|  | RNC1-510<br>(EMD-53553)<br>(PDB 9R3A) |
| **Data collection and processing** |  |
| Magnification | 75,000 |
| Voltage (kV) | 300 |
| Electron exposure (e-/Å²) | 29.8 |
| Defocus range (µm) | -1.3 to -2.8 |
| Pixel size (Å) | 1.08 |
| Symmetry imposed | C1 |
| Initial particle images (no.) | 835,178 |
| Final particle images (no.) | 389,472 |
| Map resolution (Å) | 2.86 |
| FSC threshold | 0.143 |
| Map resolution range (Å) | 2.4-8.2 |
| **Refinement** |  |
| Initial model used (PDB code) | 7K00, 8QOA |
| Model resolution (Å) | 2.86 |
| FSC threshold | 0.143 |
| Model resolution range (Å) | 2.4-8.2 |
| Map sharpening B factor (Å²) | -61.45 |
| Model composition |  |
| Non-hydrogen atoms | 235,111 |
| Protein residues | 5,625 |
| Nucleotide residues | 4,540 |
| Ligands | K:131<br>Zn:2<br>Mg:287<br>IAS:1 |
| B factors (Å²) |  |
| Protein | -103.48 |
| Ligand | -73.37 |
| R.m.s. deviations |  |
| Bond lengths (Å) | 0.006 |
| Bond angles (°) | 0.597 |
| Validation |  |
| MolProbity score | 1.22 |
| Clashscore | 2.60 |
| Poor rotamers (%) | 0.04 |
| Ramachandran plot |  |
| Favored (%) | 97.03 |
| Allowed (%) | 2.97 |
| Disallowed (%) | 0.00 |

